# Solvation Shapes the Conformational Landscape of a Therapeutically Relevant SMN2 Splice-Site Defect

**DOI:** 10.64898/2026.07.01.735918

**Authors:** Mohammed Khaled, Lucas Leuschner, Oscar Palomino-Hernandez

**Affiliations:** Department of Chemistry, Johannes Gutenberg University Mainz, Duesbergweg 10-14, D-55128 Mainz, Germany; Institute for Quantitative and Computational Biosciences (IQCB), Johannes Gutenberg University Mainz, Johannes-von-Müller-Weg 6, D-55128 Mainz, Germany

**Keywords:** SMN2, RNA simulations, molecular dynamics, replica exchange, solvation models

## Abstract

The SMN2 exon 7 5’ splice-site/U1 snRNA duplex contains an A_−1_ bulge that weakens splice-site recognition and represents a therapeutically relevant RNA connectivity defect, yet its conformational landscape and coupling to solvation remain poorly understood. Here, we performed enhanced-sampling Hamiltonian replica-exchange molecular dynamics simulations of the SMN2 splice-site duplex using four explicit-solvent models (OPC, TIP4P-Ew, TIP3P, and SPC/E) and characterized the sampled ensemble using linear and machine-learned latent representations.

Across representations, the A_−1_ defect consistently populated three metastable conformational states distinguished by local duplex geometry, base stacking, hydrogen-bonding patterns, and solvent exposure. The relative populations of these states, together with first-shell hydration and Na^+^ distributions around the defect, varied substantially across water models, demonstrating that hydration and ion organization actively shape the equilibrium between locally accommodated and solvent-exposed conformations of the SMN2 splice-site bulge.

Our results shed light on the conformational components of this therapeutic RNA target and highlight the impact of solvation model as an important consideration for molecular simulations of RNA splice-site recognition and small-molecule repair.

## Introduction

A central feature of ribonucleic acid (RNA) architecture is the presence of local deviations from ideal A-form helicity [1–3]. These structural elements include bulges, loops, mismatches, and other higher-order motifs [4] that modulate helical continuity, and allow the exposure of specific nucleobases. In this way, these elements can act as functional topological defects [5, 6], locally reshaping the RNA duplex and generating binding-competent conformations [7, 8].

Among these motifs, bulges and internal loops are particularly important because they introduce interruptions into otherwise helical regions. A bulge arises when one or more nucleotides remain unpaired on one strand of a duplex, whereas an internal loop contains unpaired nucleotides on both strands [9]. Their functional importance is therefore closely linked to their ability to remodel local RNA topology while preserving sufficient duplex character to maintain global structural coherence [4]. This dual role, both as a structural perturbation and a recognition element, is especially relevant in pre-messenger RNA (mRNA) splicing, where short RNA sequence elements must be interpreted by the spliceosome with high fidelity. Recognition of the 5’ splice site (SS) is one of the earliest steps of spliceosome assembly and involves base pairing between the 5’ end of U1 snRNA and the 5’ SS of the pre-mRNA [10]. Thus, local structural irregularities at splice sites can become molecular determinants of splice-site strength and spliceosome recruitment [11].

A prominent example of such a regulatory defect is the A_−1_ bulge located at the 5’ SS of human Survival of Motor Neuron (SMN) pre-mRNA [12]. This bulged adenosine occurs at the exon-intron junction of SMN2 exon 7 within the duplex formed between the SMN2 5’ SS and U1 snRNA (Figure 1) [13]. The presence of the A_−1_ bulge creates a local discontinuity in the RNA duplex and contributes to the weakness of the SMN2 exon 7 5’ SS [12, 13]. Because efficient U1 snRNP recruitment is required for productive spliceosome assembly, this local RNA structural defect is directly connected to inefficient SMN2 exon 7 inclusion [13–15].

**Figure 1:**
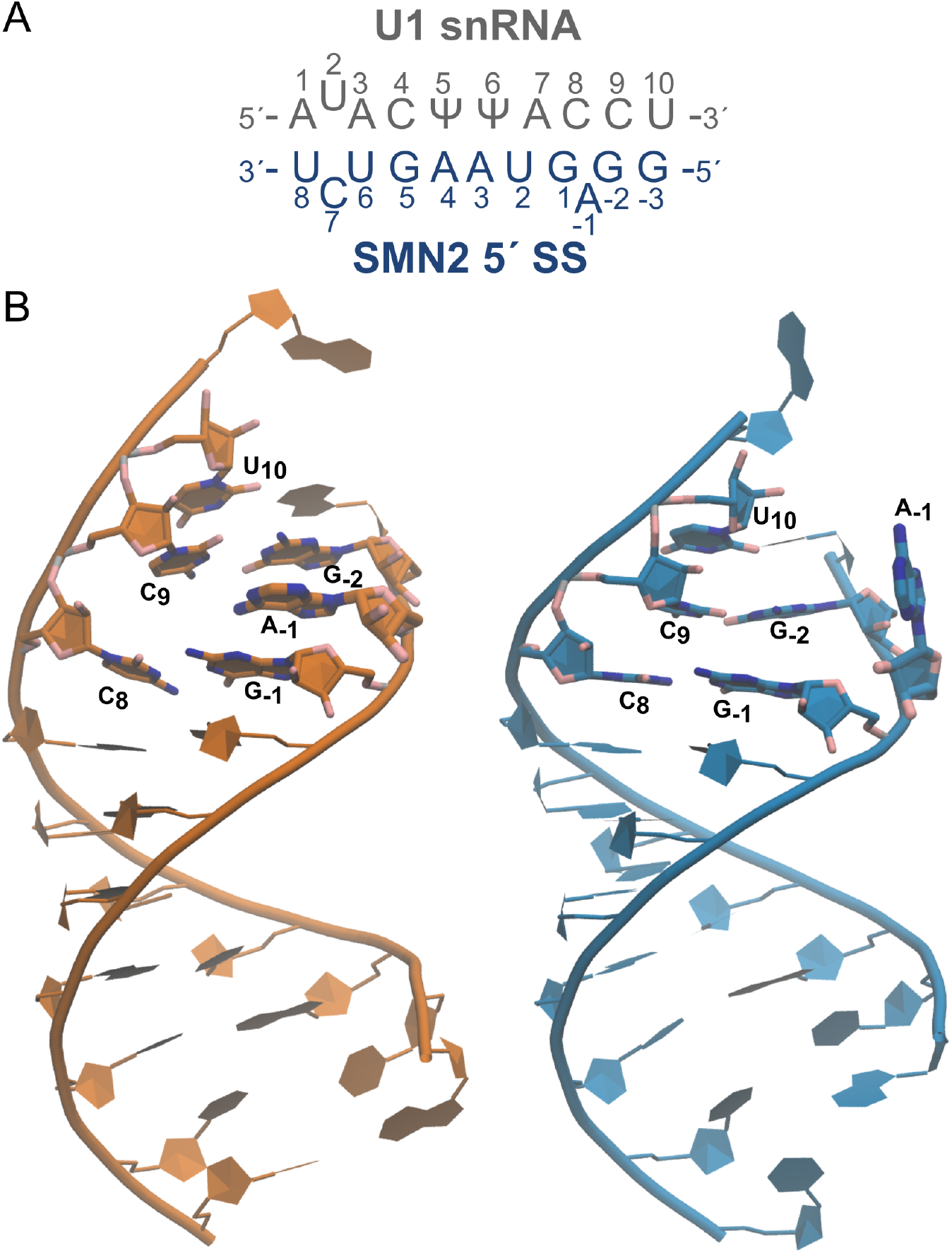
(A) Schematic diagram showing the RNA-RNA duplex formed between U1 snRNA and and the SMN2 5’ SS. (B) Representative solution structures from the experimental NMR 6HMI (left) and 6HMO (right), with the A_−1_ bulge region nucleotides C_8_, C_9_, U_10_, G_1_, A_−1_ and G_−2_ highlighted in stick representation.

The biological importance of this splice-site defect is highlighted by the genetic context of Spinal Muscular Atrophy (SMA). SMA is primarily caused by deletion or mutation of Survival of Motor Neuron 1 (SMN), which reduces the production of functional SMN protein [16]. Humans also possess the paralogous Survival of Motor Neuron 2 (SMN) gene, but SMN2 cannot fully compensate for loss of SMN1 because most SMN2 transcripts skip exon 7 and produce an unstable protein isoform [17]. Restoration of SMN2 exon 7 inclusion through A_−1_ bulge stabilization is therefore a major therapeutic strategy, as increased production of full-length SMN protein can partially compensate for SMN1 deficiency [13, 14]. Structural studies of small-molecule splicing modifiers have shown that compounds such as SMN-C5 bind near the exon-intron junction, stabilize the bulged adenine, and convert the weak SMN2 exon 7 5’ SS into a stronger splice-site substrate [13]. More recent work further indicates that chemically diverse splicing modifiers can act on related A_−1_-bulged 5’ SS architectures, supporting the broader concept that specific RNA connectivity defects can be pharmacologically leveraged [12, 15, 18].

Consequently, the intrinsic conformational landscape of the SMN2 RNA splice-site defect remains an important unresolved question. In particular, it is unclear how the A_−1_ bulge behaves in the conformational ensemble, and how the local RNA environment modulates the energetic accessibility of repaired versus unrepaired states [13, 15, 18]. Addressing these questions requires a dynamic view of the defect [19]. For this, we performed enhanced-sampling replica-exchange molecular dynamics (MD) simulations of the SMN2 splice-site RNA system. We explored the conformational propensities of the A_−1_ bulge, the stability of local base-pairing interactions, and the metastable states that connect disrupted and repaired duplex states. Furthermore, we explored the role of solvation in the system, as it has been shown to be a critical but often overlooked factor [20]. By characterizing these ensembles, we aim to clarify how local RNA topology and its environmental in solution shape the conformational accessibility of the SMN2 splice-site defect. This dynamic perspective provides a framework for understanding how stabilization of the A_−1_ bulge can contribute to restoration of SMN2 exon 7 inclusion.

## Results

### The SMN2 A_−1_ Defect Populates Three Conformational States

We first aimed to characterize the conformational landscape of the A_−1_ defect in the RNA duplex formed with U1 snRNA. To this end, enhanced-sampling Hamiltonian replica-exchange simulations of the duplex were performed for every system for at least 3.0 *μ*s of cumulative sampling time. To evaluate the influence of the solvent environment, simulations were carried out using four explicit-solvent models: OPC, TIP4P-Ew, TIP3P, and SPC/E.

To describe the conformational landscape, structures were extracted from each trajectory, concatenated, and represented using a 22-dimensional feature space. These descriptors capture the structural organization of the A_−1_ bulge and its local environment, including local geometry, base-pairing interactions, angles, dihedrals, and solvent exposure within the C_9_-A_−1_ region (Supplementary Figure S1). This representation provides an invariant description of the local RNA defect.

Because the resulting conformational space is intrinsically high-dimensional, we applied dimensionality-reduction with principal component analysis (PCA) to identify the dominant metastable states [21]. In parallel, multiple non-linear encoder-decoder neural network models were trained in order to use their latent spaces as alternative reduced-dimensional representations. We explored the compression capability of each model by varying the output dimension of the encoder network between *d ∈* 2, 4, 6, 8. These models included Autoencoder (AE), Denoising Autoencoder (DAE), Contracting Autoencoder (CAE), and variational architectures such as Variational Autoencoder (VAE), *β*-Variational Autoencoder (BVAE), Variational Autoencoder with laplacian prior (VLAE) and Dirichlet Variational Autoencoder (DVAE). Further information regarding the hyperparameter optimization of these models is provided in the Supporting Information (Supplementary Tables S1-S4). To identify metastable states, HDBSCAN clustering was applied to each reduced-dimensional representation [22]. The combined use of linear and non-linear approaches allowed us to evaluate whether the inferred latent space was robust to the choice of projection method and dimensionality.

Under PCA, projection of the conformations onto the first two principal components retained approximately 56% of the total variance. This initial representation already revealed multiple metastable basins corresponding to distinct structural states of the A_−1_ bulge (Supplementary Figure S2). To capture a larger fraction of the conformational variance, the PCA representation was then expanded to include more components, recovering ∼75% at six dimensions and up to ∼83% at eight dimensions (Supplementary Table S5). In this higher-dimensional PCA space, HDBSCAN resolved three conformational states together with a sparsely populated heterogeneous ensemble classified as noise (Figure 2).

**Figure 2:**
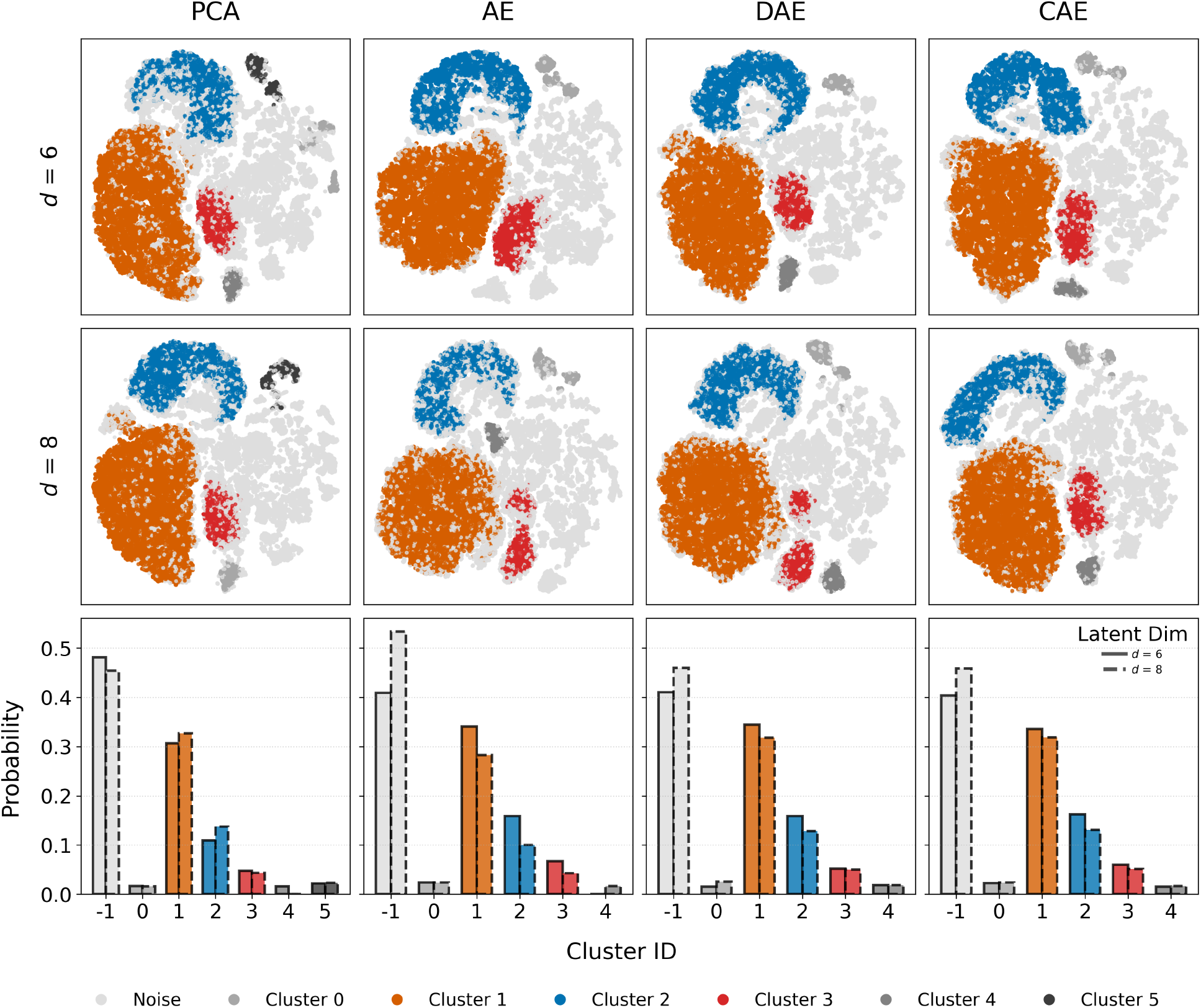
Two-dimensional projections of the conformational space generated via *t*-SNE applied to the latent spaces (*d* = 6 and *d* = 8) of PCA, AE, DAE, and CAE. Cluster probabilities extracted across the corresponding embedding dimensions (*d* = 6, 8). Clusters are color-coded as follows: Cluster 1 (orange), Cluster 2 (blue), and Cluster 3 (red), while noise clusters are shown in shades of gray.

The non-linear models provided an independent view of this state decomposition. Deterministic architectures focused on reconstruction accuracy (such as AE, DAE, and CAE) reproduced the metastable states identified by PCA more consistently than the variational models (Supplementary Figure S3). Among them, the DAE model was particularly effective, preserving the cluster topology with high fidelity at latent dimensions of *d* = 6 and *d* = 8 (Figure 2). In contrast, variational approaches generated smoother latent manifolds that reduced the separation between conformational basins, leading to increased mixing between states during HDBSCAN analysis (Supplementary Figure S3). This behavior is consistent with the regularizing effect of the variational prior, which promotes continuity in the latent space at the cost of compressing distinct conformational basins into less separated regions.

Across all architectures, lower latent dimensions (*d* = 2 and *d* = 4) were insufficient to represent the system accurately. These compressed embeddings often resulted in over-segmentation of the data into numerous small clusters or unstable assignment of conformations to metastable states (Supplementary Figure S4). This indicates that the conformational manifold of the A_−1_ bulge cannot be represented reliably in very low-dimensional embeddings and instead requires a moderately high-dimensional latent space to preserve the relevant structural organization.

Across the different dimensionality-reduction methods and latent-space dimensions, the cluster population analysis showed a consistent dominance of two major states, assigned as clusters 1 and 2 (Table 1). These states remained stable across PCA and deterministic autoencoders, particularly for *d* = 6 and *d* = 8. In contrast, Cluster 3 appeared as a lower-population but reproducible metastable state whose occupancy varied more strongly with the embedding and clustering procedure. Other clusters were not consistently recovered across models and were therefore interpreted as method-dependent fragmentation of the conformational space rather than independent major states (Figure 2 and Supplementary Figures S3 and S4). Although variational models further reduced cluster separability, they did not alter the overall ranking of the dominant basins.

**Table 1:**
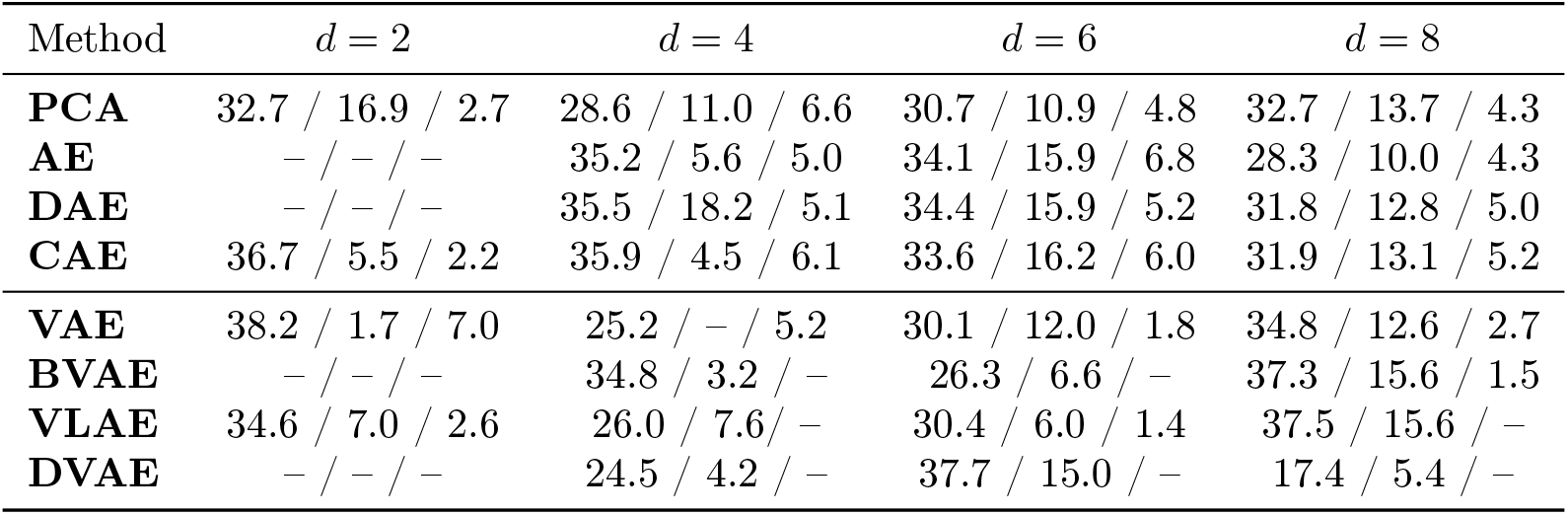
Cluster population distributions (%) obtained from PCA and different autoencoder-based embeddings at varying latent dimensions (*d* = 2, 4, 6, 8). Values correspond to Cluster 1–3 for each method and embedding size.

Taken together, these analyses indicate that the SMN2 A_−1_ defect organizes into a small number of recurrent metastable conformational states that capture the essential structure of the SMN2 A_−1_ conformational equilibrium.

### Structural Characterization of the Conformational States

Having established that the SMN2 A_−1_ defect populates three recurrent metastable states, we next characterized the structural features that distinguish these conformational basins. For each state, representative structures were extracted and compared in terms of geometry, solvent exposure of the bulged adenine, hydrogen-bonding patterns, and local stacking interactions, in order to understand the role each major component plays in the conformational ensemble.

The structural characterization of the three identified clusters reveals distinct interaction networks and conformational preferences (Figure 3). Cluster 1 most closely resembles the experimentally observed ligand-bound structure (PDB: 6HMO, Supplementary Figure S5A), in which the A_−1_ nucleotide is locally accommodated within the duplex and the neighboring base pairs preserve a continuous helical arrangement. Cluster 2 most closely resembles the experimentally observed *apo* structure (PDB: 6HMI, Supplementary Figure S5B), in which the bulged nucleotide remained partially associated with the duplex but did not fully reproduce the repaired-like geometry observed in the experimental structure. In contrast, Cluster 3 showed a more distorted geometry, characterized by partial disruption of the local stacking network.

**Figure 3:**
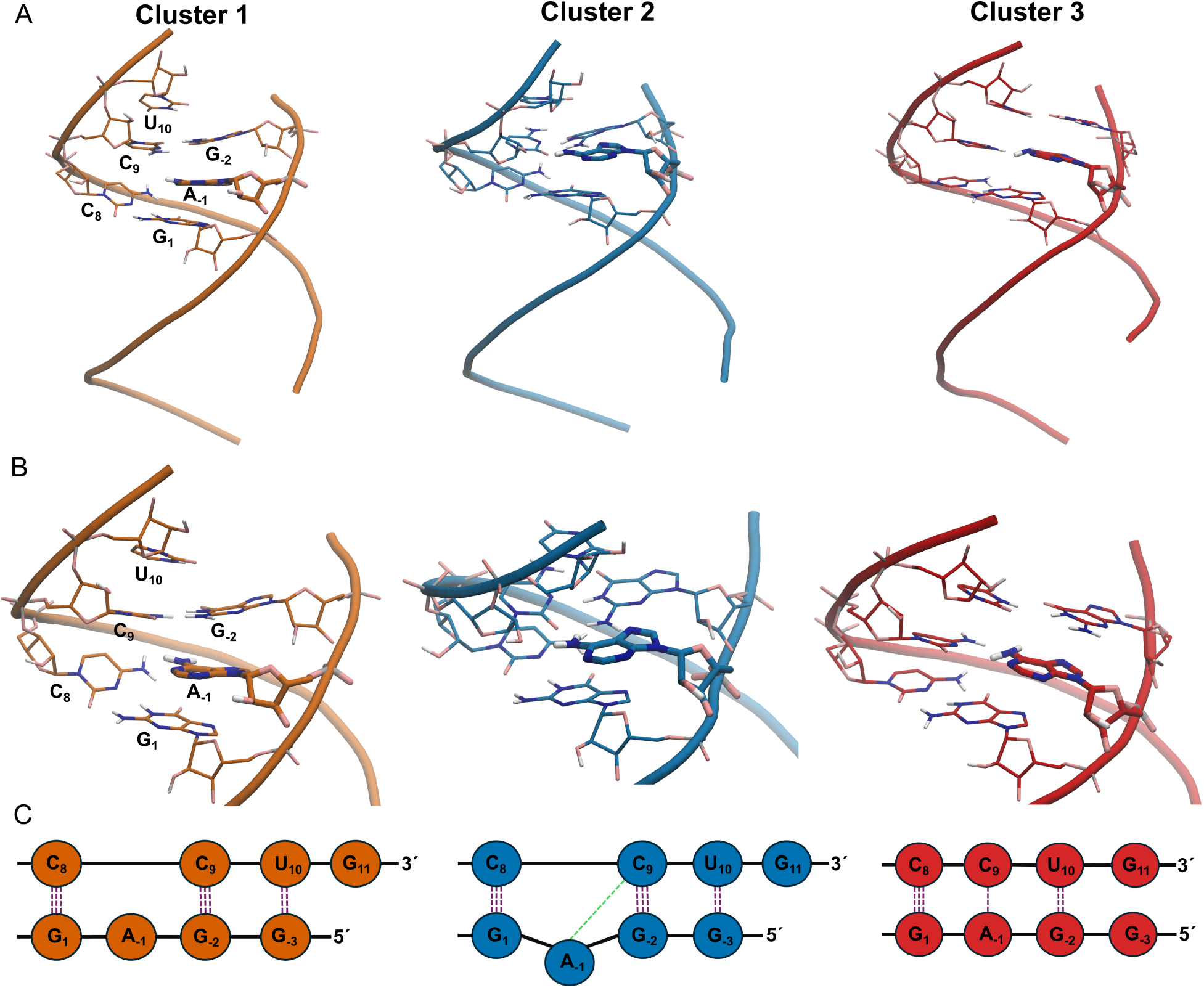
(A) Representative structures of the three clusters are shown, with nucleotides C_8_, C_9_, U_10_, G_1_, A_−1_ and G_−2_ highlighted as sticks. (B) A zoomed-in view of the A_−1_ bulge region. (C) Schematic interaction networks illustrating the nucleotide base topologies surrounding the A_−1_ bulge. Green dashed lines indicate Sugar-Hoogsteen cis (SHc) interaction, while purple dashed lines indicate Watson-Crick cis (WWc) interactions; the number of the lines corresponds to the average number of hydrogen bonds.

To further characterize these states, we analyzed the hydrogen-bonding networks between A_−1_ and its neighboring residues (Figure 4), RNA base-pair interactions (Table 2 and Supplementary Table S7), consecutive stacking interactions (Table 3 and Supplementary Table S8), and residue-specific solvent-accessible surface area (SASA) values within the A_−1_ region (Figure 4). We observed that the three conformational states are distinguished by different balances between canonical base pairing, non-canonical hydrogen bonding, vertical stacking, and solvent exposure.

**Table 2:**
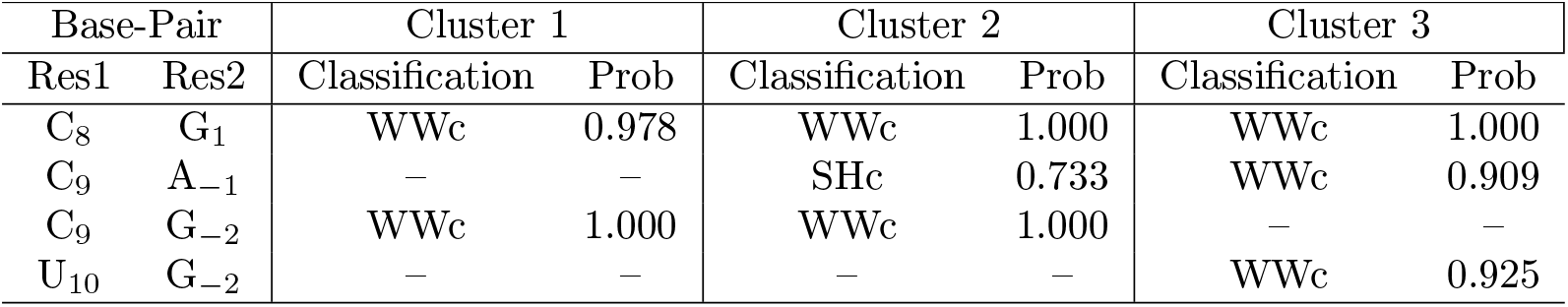
Base-pair interactions of A_−1_ surrounding residue and their probabilities across the three clusters. Base pairs are classified according to the Leontis-Westhof scheme: W, H, and S denote Watson-Crick, Hoogsteen, and Sugar edges, respectively, while c/t indicate cis or trans orientation.

**Table 3:**
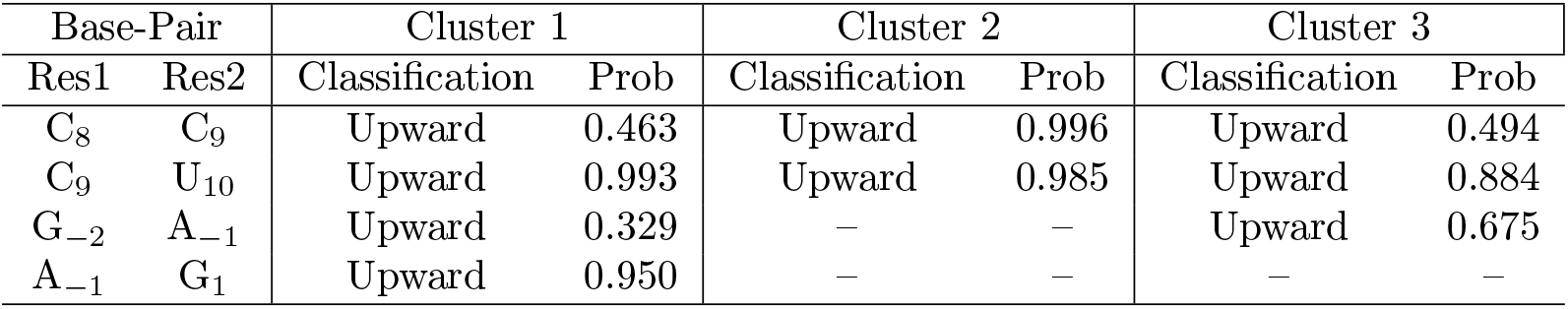
Consecutive Base-pair stacking interactions of A_−1_ surrounding residue and their probabilities across the three clusters. Base pairs are classified as upward (3’-5’), downward (5’-3’), outward (5’-5’), and inward (3’-3’).

**Figure 4:**
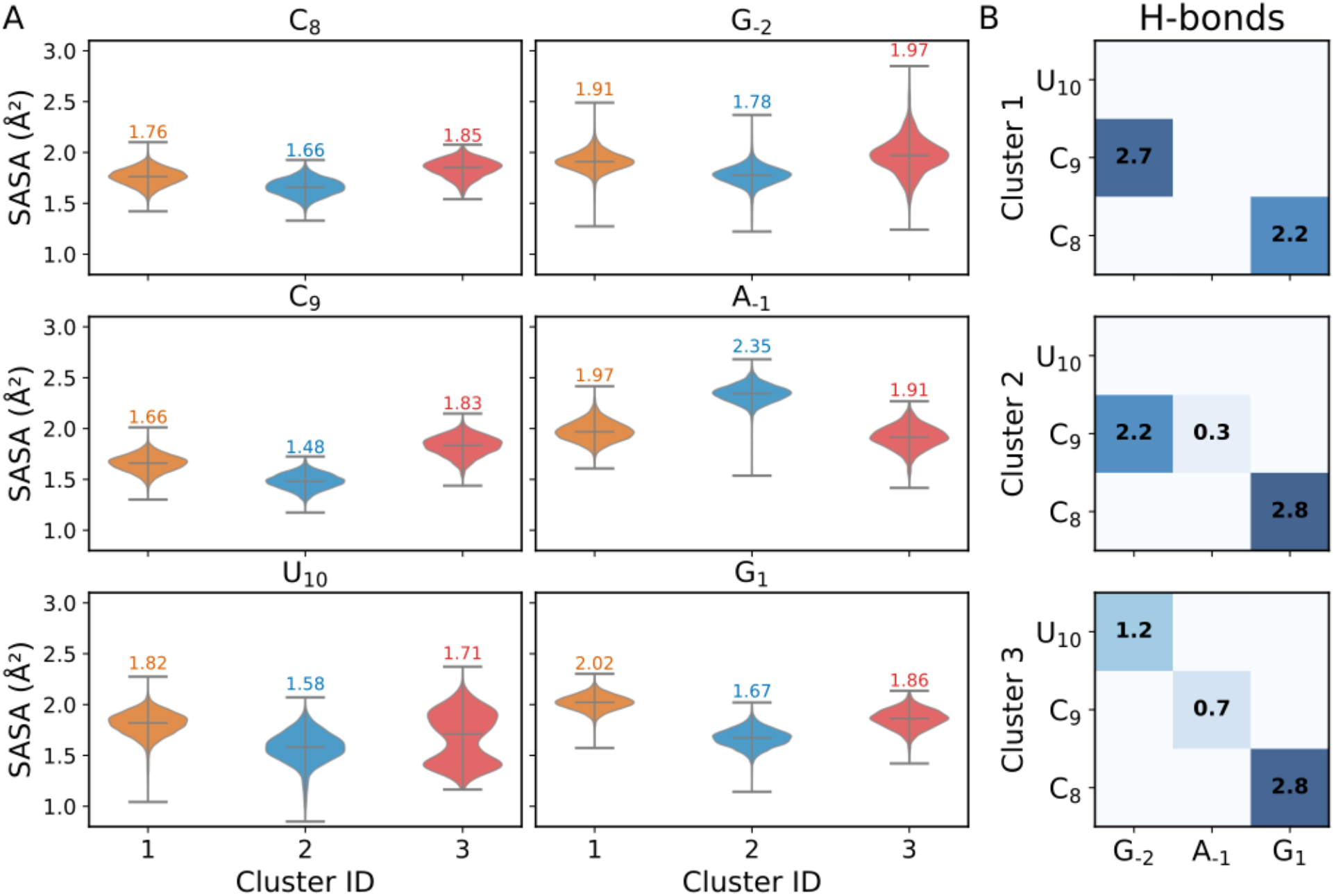
A) Distribution of solvent-accessible surface area (SASA) for C_8_, C_9_, U_10_, G_1_, A_−1_ and G_−2_ across the clusters. The text labels indicate the median SASA values. B) Inter-residue hydrogen bonding networks across the clusters. The heatmaps display the mean number of hydrogen bonds between the G_1_-A_−1_-G_−2_ and C_8_-C_9_-U_10_ residue groups for each water model. Numerical labels indicate the average hydrogen bond number.

Cluster 1 (stacked-unbound state) is characterized by preservation of the canonical C_8_-G_1_ and C_9_-G_−2_

Watson-Crick base pairs together with persistent stacking interactions between A_−1_ and the neighboring G_1_ and G_−2_ bases (Tables 2 and 3). In this configuration, A_−1_ remains integrated within the helical core despite the absence of stable hydrogen-bonding interactions involving the adenine base itself. Consistently, A_−1_ shows relatively low SASA, indicating that the bulged nucleotide is partially sequestered from the solvent (Figure 4). Thus, the stability of this state is primarily associated with vertical stacking rather than base pairing. This arrangement maintains a compact, A-form-like local geometry, also reflected by an increase in duplex inclination relative to the helical axis (Supplementary Figure S6).

Cluster 2 (solvent-exposed state) is defined by a marked reorganization of the A_−1_ bulge. In this state, A_−1_ leaves the previous stacking towards a more solvent-exposed structure (Figure 4). This state is stabilized by a transient hydrogen bond in which the N6 amino group of A_−1_ acts as a donor to the carbonyl oxygen O2 of C_9_ (Figure 4). This effectively produces a non-canonical triplet-like arrangement in which C_9_-G_−2_ maintains a Watson–Crick cis (WWc) interaction, although weakened to two persistent hydrogen bonds, while A_−1_ interacts with C_9_ through a Sugar–Hoogsteen cis (SHc) interface (Table 3).

This local rearrangement also affects the surrounding duplex geometry. The inclination of the base pairs relative to the helical axis decreases (Supplementary Figure S6), indicating a loss of local compactness. At the same time, although A_−1_ becomes more solvent exposed, the total RNA SASA decreases relative to the most open conformational state (Supplementary Figure S5C). This apparent contrast can be explained by the reciprocal exposure profile observed for C_9_: in Cluster 2, C_9_ becomes partially shielded from solvent by its retained interaction with G_−2_ and by the in-plane capping interaction provided by the shifted A_−1_ residue. Thus, Cluster 2 is locally open with respect to A_−1_, but partially compacted around C_9_.

Cluster 3 (registry-shift state) is defined by a deeper reorganization of the local hydrogen-bonding network and corresponds to the most globally open metastable configuration. In this state, the interaction pattern around the defect is displaced by two non-canonical base-pairing interactions: C_9_-A_−1_ and U_10_-G_−2_, both classified as Watson-Crick cis interactions (Table 2). Formation of the C_9_-A_−1_ interaction requires a lateral shift of A_−1_ and the loss of vertical stacking between A_−1_ and G_1_, thereby disrupting the local helical packing around the bulge (Table 3). This rearrangement is reflected in the changes of O2^′^-O2^′^ inter-sugar distances for the C_9_-G_−2_ and C_9_-A_−1_ residue pairs (Supplementary Figure S7), indicating a significant departure from canonical A-form geometry. Given the role of the 2^′^-hydroxyl group in RNA backbone flexibility and conformational sampling [23], these increased distances suggest a more strained local sugar-phosphate environment. The displacement of U_10_ and G_−2_ from their standard positions further weakens local duplex integrity in Cluster 3. As a result, this state exhibits the highest total RNA SASA across the three conformational states (Supplementary Figure S5C).

Together, these results demonstrate that the conformational variability of the SMN2 A_−1_ bulge lies in an equilibrium between base stacking, non-canonical hydrogen bonding, and solvent exposure. Cluster 1 is primarily stabilized by stacking and retains a compact duplex-like arrangement. Cluster 2 is defined by solvent exposure of A_−1_ and formation of a triplet-like A_−1_-C_9_-G_−2_ geometry. Cluster 3 corresponds to a state stabilized by alternative non-canonical base pairing.

### The Conformational Equilibrium of the A_−1_ Defect Is Strongly Dependent on the Water Model

We next examined how the populations of the conformational states depend on the explicit-solvent model used in the simulations. Since the A_−1_ nucleotide can alternate between buried, partially stacked, and solvent-exposed arrangements, differences in water-model behavior may directly influence the stability of repaired-like and disrupted conformations. To evaluate this effect, the cluster assignments obtained from the reduced-dimensional representations were projected back onto the unbiased MD trajectories generated for each water model. This allowed us to compare the relative populations of the metastable states across OPC, TIP4P-Ew, TIP3P, and SPC/E simulations.

Although all water models sampled the principal conformational states identified above, substantial differences were observed in their relative populations (Figure 5 and Table 4). OPC, TIP4P-Ew, and SPC/E exhibited a conserved preference for Cluster 1, with relative populations ranging from 71 to 75%. In contrast, TIP3P shifted the equilibrium toward Cluster 2, which became its most populated state with an occupancy of 64%. In addition, OPC and TIP3P displayed an increased preference for Cluster 3 in comparison to other water models.

**Table 4:**
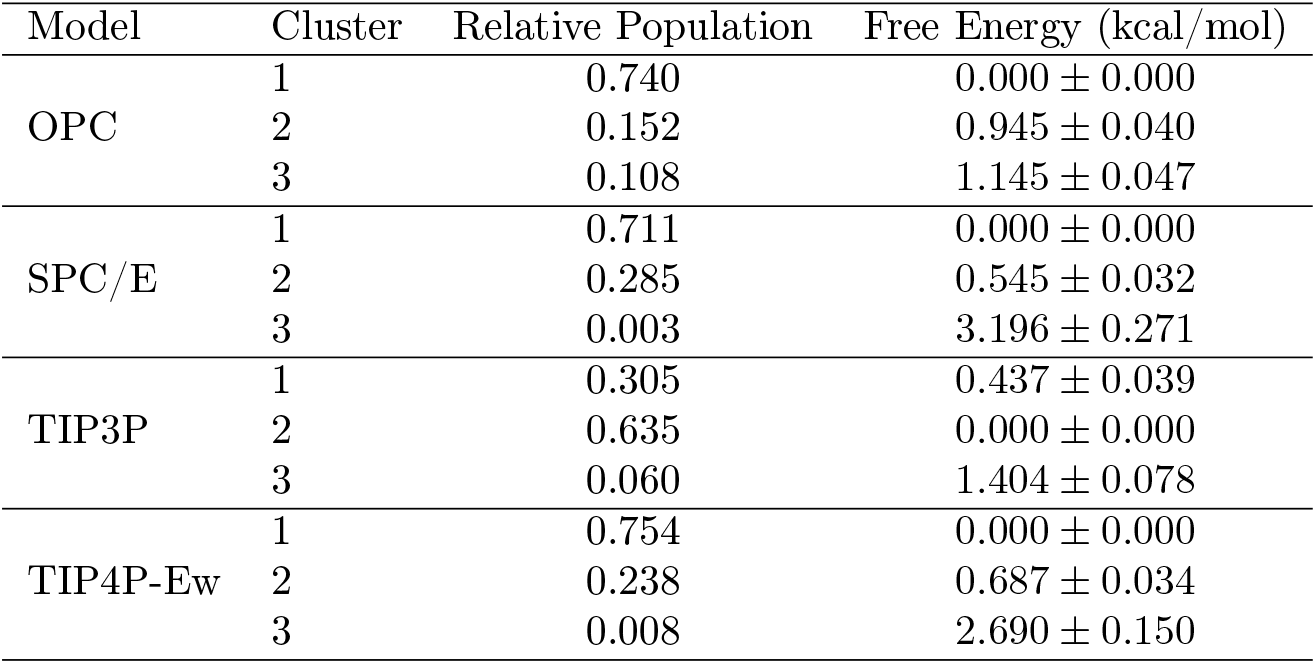
Relative populations and free energies of the three clusters for each water model. Free energies are reported relative to the most populated cluster within each model and are given as mean and bootstrap standard deviation computed over 50 bootstrap samples.

| Model | Cluster | Relative Population | Free Energy (kcal/mol) |
| --- | --- | --- | --- |
| OPC | 1 | 0.740 | 0.000 ± 0.000 |
| | 2 | 0.152 | 0.945 ± 0.040 |
| | 3 | 0.108 | 1.145 ± 0.047 |
| SPC/E | 1 | 0.711 | 0.000 ± 0.000 |
| | 2 | 0.285 | 0.545 ± 0.032 |
| | 3 | 0.003 | 3.196 ± 0.271 |
| TIP3P | 1 | 0.305 | 0.437 ± 0.039 |
| | 2 | 0.635 | 0.000 ± 0.000 |
| | 3 | 0.060 | 1.404 ± 0.078 |
| TIP4P-Ew | 1 | 0.754 | 0.000 ± 0.000 |
| | 2 | 0.238 | 0.687 ± 0.034 |
| | 3 | 0.008 | 2.690 ± 0.150 |

**Figure 5:**
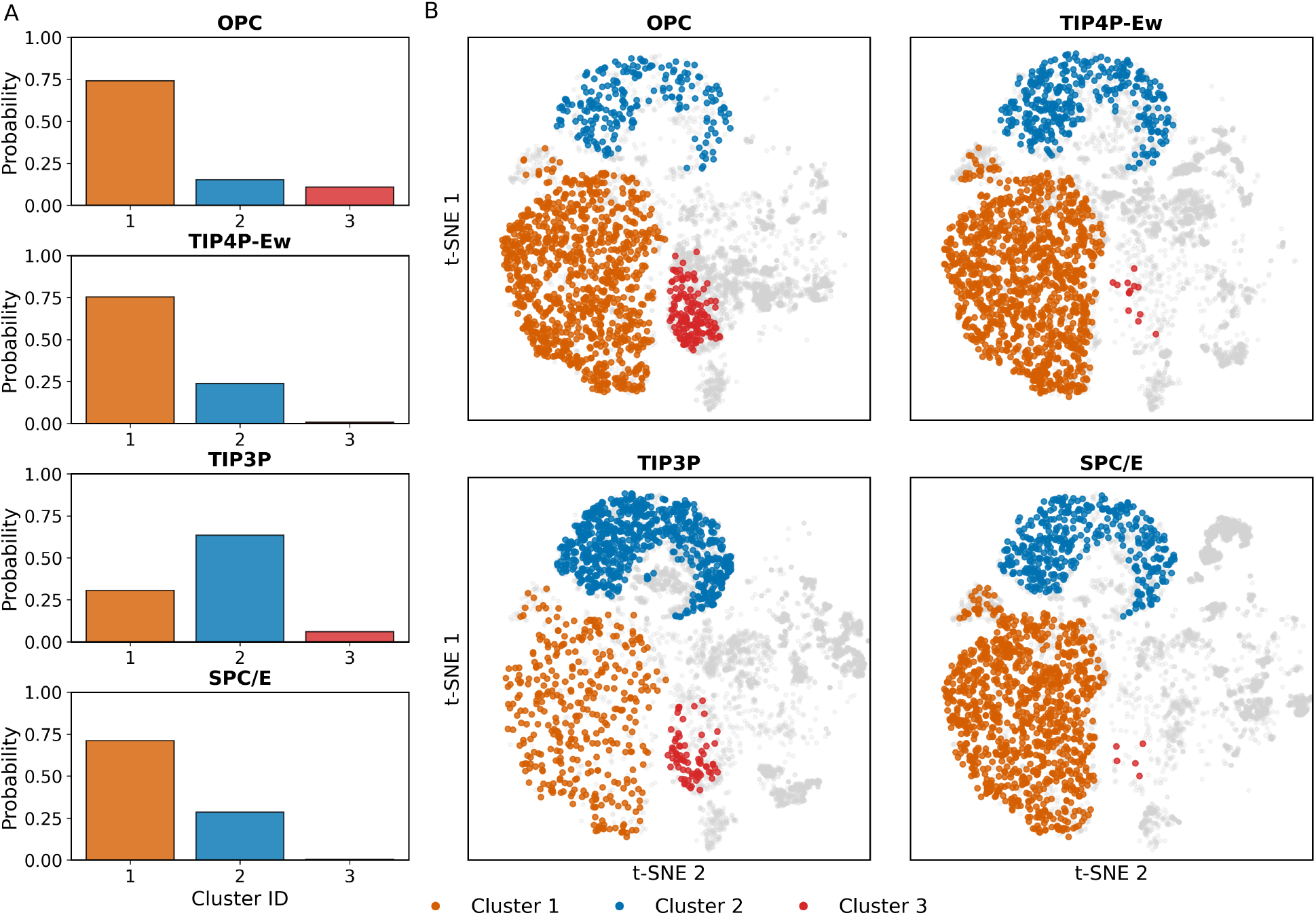
A) Relative cluster probabilities for different water models. B) Two-dimensional projection of the conformational space obtained using t-SNE applied to the first eight principal components for OPC, TIP4P-Ew, TIP3P, and SPC/E water models. Clusters are color-coded as follows: Cluster 1 (orange), Cluster 2 (blue), and Cluster 3 (red), while noise points are shown in light gray.

Thus, we notice a strong change depending on the water model: OPC, TIP4P-Ew, and SPC/E show a preference for stacking interactions, where the A_−1_ defect is located inside the A-helix. In contrast, TIP3P seems to stabilize the solvent-exposed state, either via favorable A_−1_-solvent interactions, from a reduced stabilization of local base-stacking interactions, or from a combination of both effects. Overall, it is remarkable that there is such a strong solvent-model dependence of the SMN2 A_−1_ conformational equilibrium. This observation is particularly relevant for interpreting small-molecule repair of the SMN2 5’ SS, since we would expect the structures where A_−1_ is solvent-exposed to be the lower energy ones. Thus, for this system, the choice of water model inverted the relative stability of metastable states.

### Atomistic Insights into Solvent and Ion Organization around the A_−1_ Defect

The strong dependence of the conformational equilibrium on the water model suggests that solvation plays an active role in shaping the local RNA defect landscape. Water molecules contribute to dielectric screening, stabilize exposed polar groups, and modulate the balance between base stacking and solvent exposure. In addition, RNA is surrounded by a counterion atmosphere that shields the negative charge of the phosphate backbone [24]. These counterions associate with RNA at different degrees of hydration and often interact indirectly through coordinated water molecules within the diffuse ion atmosphere [25]. This dynamic solvent-ion environment can rapidly reorganize in response to local conformational fluctuations, thereby contributing to the structural plasticity of the RNA. Such effects are particularly relevant for a bulged nucleotide such as A_−1_, whose conformational states differ in the extent to which the base and backbone are buried within the duplex or exposed to surrounding water molecules and ions.

To characterize the local solvent organization around the defect, we analyzed the spatial distribution of water molecules and Na^+^ ions around the RNA residues across the identified conformational clusters. We first quantified the average occupancy of water molecules and ions within the first solvation shell of the phosphate groups and then complemented this analysis with radial distribution functions (RDFs) around the A_−1_ residue. In addition, three-dimensional density maps of water and sodium ions were calculated to visualize preferential ion and water accumulation around the RNA duplex (Supplementary Figures S8-S9) and around A_−1_ residue (Figure 8).

When focusing on the first hydration shell, defined using a cutoff of 3.5 Å, we observed pronounced differences in the local organization of water molecules around the RNA phosphate backbone (Figure 6A). Although most water models showed comparable hydration profiles, TIP3P consistently exhibited higher water occupancy near the phosphate groups. This enrichment was observed across all clusters, indicating a systematic increase in local phosphate hydration. In contrast, OPC generally showed lower water occupancy than the other solvent models, suggesting a reduced tendency to populate the immediate hydration shell of the RNA backbone.

**Figure 6:**
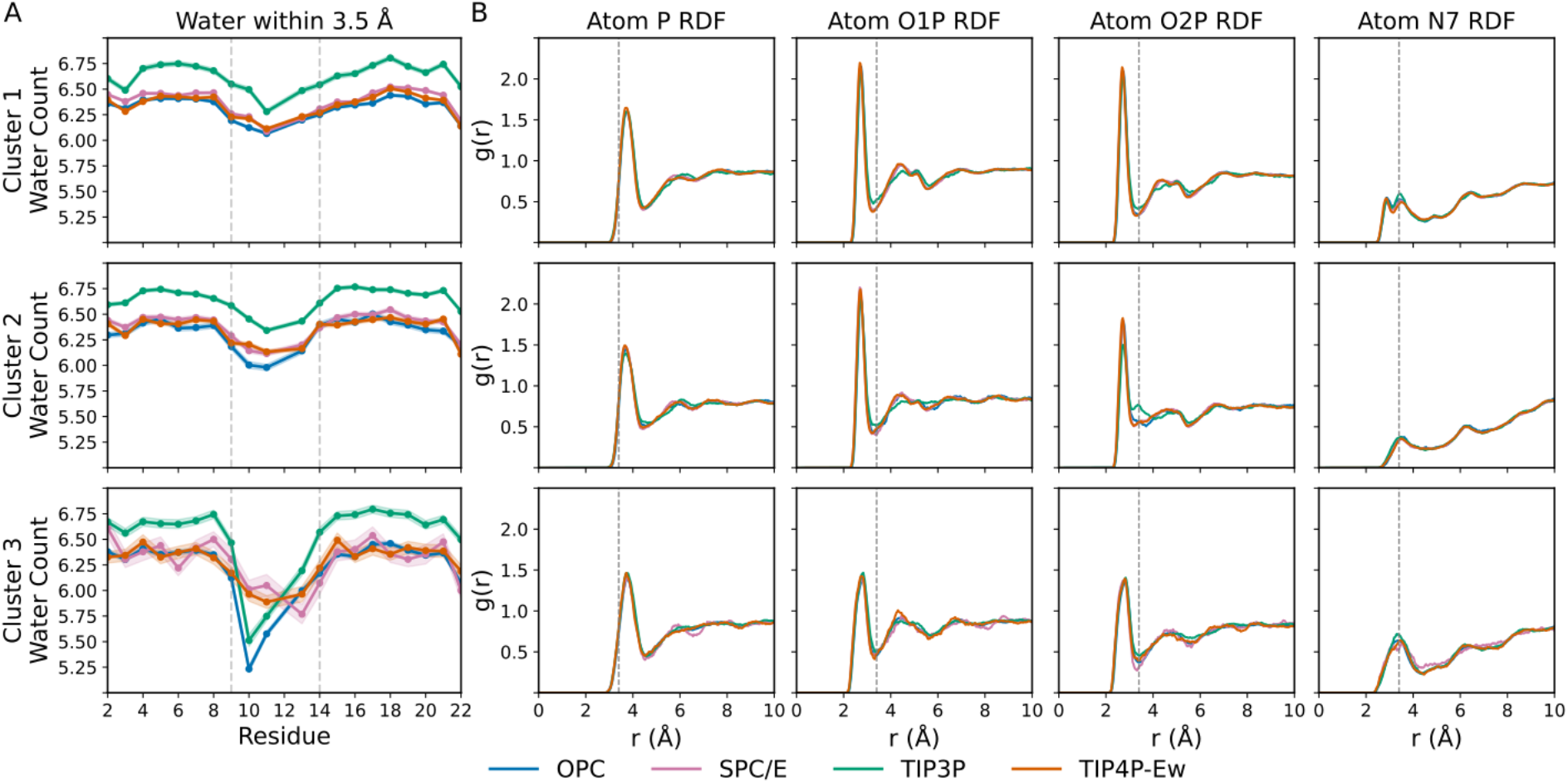
(A) Average occupancy of water within 3.5 Å of the phosphate group atpms (P, O1P, and O2P) per residue across the clusters. The shaded regions denote the Standard Error of the Mean (SEM). Vertical dashed lines highlight the positions of the C_9_ and A_−1_ residues. (B) Radial distribution functions (RDFs) of waters around the A_−1_ phosphate group (P, O1P and O2P) and N7 atom. The plots show the average RDF, *g*(*r*), for the three Clusters across the four water models. A dashed black vertical line at 3.4 Åindicates the boundary of the first solvation shell. Water models are color-coded as follows: OPC (green), TIP4P-Ew (orange), TIP3P (blue), and SPC/E (purple).

A more local comparison revealed marked changes around residues U_10_ and G_11_, which define the region adjacent to the A_−1_ defect. For these residues, the hydration pattern deviated from the global residue-wise trend. In particular, both OPC and TIP3P showed reduced water occupancy in Cluster 3, despite their opposite behavior in the overall hydration analysis. The reduced hydration in this region coincided with different sampling of the backbone dihedrals of these terminal residues, a behavior not observed to the same extent in the other water models (Supplementary Figure S10).

Focusing on A_−1_, the water RDFs revealed only modest differences in hydration around the phosphate backbone (Figure 6B). The largest difference was observed for the O2P atom in TIP3P, where a small additional peak indicated a more pronounced local interaction with water molecules. No strong changes were also perceived for other atoms from A_−1_ (Supplementary Figures S11-S13). This enhanced hydration of the A_−1_ phosphate group may contribute to the increased stabilization of Cluster 2 observed in TIP3P relative to the other water models.

Because the RNA backbone is highly negatively charged, Na^+^ ions are expected to accumulate near phosphate groups and screen local electrostatic repulsion. This behavior was observed in the sodium density maps (Supplementary Figure S9), but the extent of ion accumulation depended strongly on the water model (Figure 7A). Similar to the water-occupancy analysis, OPC displayed the weakest local Na^+^ enrichment around the RNA, suggesting that ions remain more diffusely distributed in the bulk solvent. In contrast, TIP3P showed the strongest association of Na^+^ ions with the phosphate backbone, particularly pronounced for Cluster 2.

**Figure 7:**
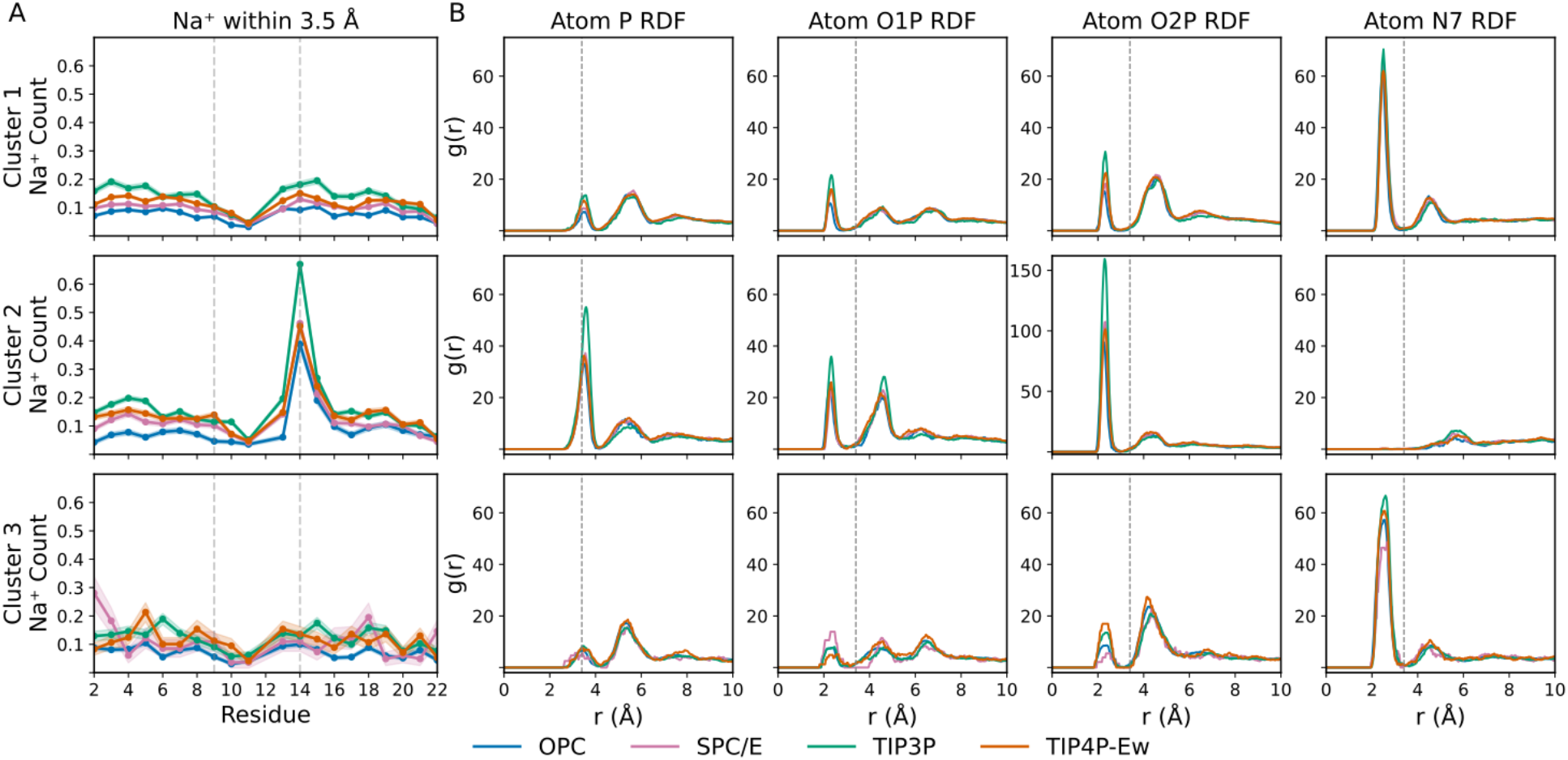
(A) Average occupancy of sodium ions (*Na*^+^) within 3.5 Å of the phosphate group atpms (P, O1P, and O2P) per residue across the clusters. The shaded regions denote the Standard Error of the Mean (SEM). Vertical dashed lines highlight the positions of the C_9_ and A_−1_ residues. (B) Radial distribution functions (RDFs) of *Na*^+^ ions around the A_−1_ phosphate group (P, O1P and O2P) and N7 atom. The plots show the average RDF, *g*(*r*), for the three Clusters across the four water models. A dashed black vertical line at 3.4 Åindicates the boundary of the first solvation shell. Note that the y-scale of atom O2P and Cluster 2 is different for clarity. Water models are color-coded as follows: OPC (green), TIP4P-Ew (orange), TIP3P (blue), and SPC/E (purple).

**Figure 8:**
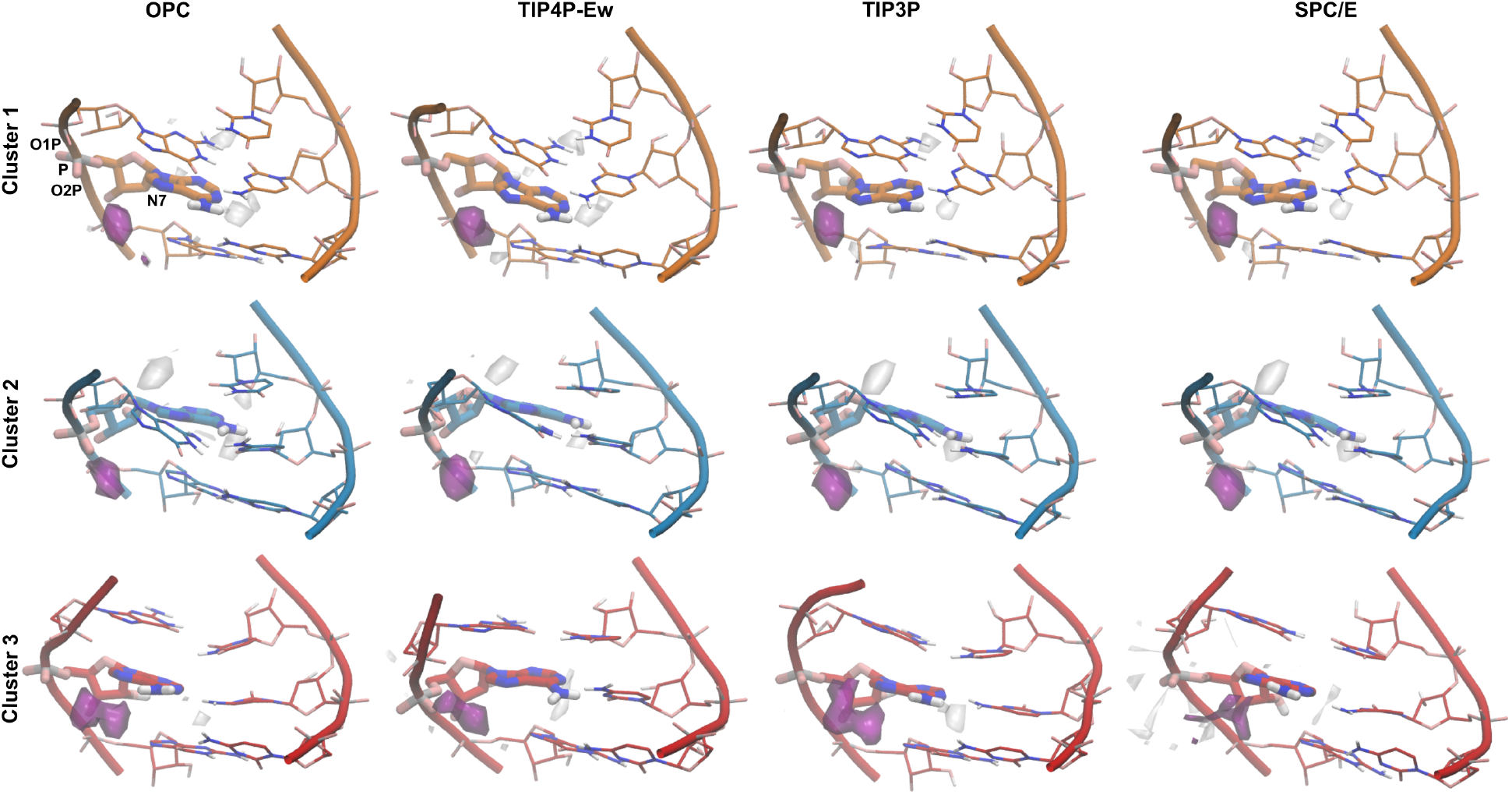
Three-dimensional density maps of sodium ions Na^+^ and water around the A_−1_ were calculated for the three clusters across the four water models (OPC, TIP4P-Ew, TIP3P, and SPC/E). The density maps were visualized using VMD and displayed at the same isocontour level of 0.009 for Na^+^ ions (0.018 for Cluster 3 of SPC/E models) and 0.06 for water. The *Na*^+^ and water density maps are shown in purple and white, respectively.

The Na^+^ RDFs also showed a marked increase in ion localization around the A_−1_ backbone atoms in TIP3P, especially in Cluster 2 (Figure 7B). In addition to phosphate coordination, Na^+^ ions also showed stronger interaction with the N7 atom of the adenine base than was observed for water molecules in the corresponding RDFs. Such interactions could help stabilize conformations in which A_−1_ is more solvent accessible, thereby linking ion localization to the conformational state of the bulge. No significant changes in Na^+^ ion coordination were observed for other atoms from A_−1_ (Supplementary Figures S14-S16).

Together, these findings indicate that the conformational equilibrium of the SMN2 A_−1_ defect is coupled to the local organization of both water molecules and Na^+^ ions. The different water models not only alter the overall hydration level of the RNA backbone, but also redistribute solvent and ions in a residue- and state-dependent manner.

## Discussion

In this work, we investigated the intrinsic conformational landscape of the SMN2 A_−1_ splice-site defect through enhanced-sampling molecular simulations and machine-learned latent representations. Our results show that the A_−1_ defect is better represented by an ensemble composed of three metastable states whose populations are strongly modulated by the solvent environment. In one state, local stacking interactions preserve duplex continuity around the exon-intron junction, resulting in a stacked conformation of the defect (Cluster 1). A second state corresponds to a disrupted arrangement, in which the bulged adenine becomes solvent-accessible and the local stacking network is weakened (Cluster 2). The third state, a minor one, resembles a repaired-like geometry, where rearranged hydrogen-bonding patterns help retain A_−1_ within the helical environment (Cluster 3). Thus, the A_−1_ bulge behaves as a functional topological defect that creates alternative structural states that can be differentially stabilized.

The identification of these states relied on the comparison of several dimensionality-reduction strategies. While PCA provided a direct view of the conformational landscape, its low-dimensional projections ignored part of the conformational heterogeneity. Deterministic autoencoders, particularly DAE models, preserved the topology of the conformational basins more effectively at latent dimensions of *d* = 6 and *d* = 8. In contrast, variational architectures produced smoother latent manifolds and reduced the separation between basins, likely because the imposed latent-space regularization favors continuity in the learned representation.

This behavior is consistent with a conformational landscape composed of discrete metastable states separated by relatively sharp structural transitions. For this system, reconstruction-focused latent spaces were better suited than variationally-regularized latent spaces to preserve the local conformational organization required for clustering. Importantly, the same major states were recovered across multiple representations, supporting the robustness of the three-state description.

The combined hydration and ion analyses provide an atomistic explanation for this solvent dependence. Stacked states (Cluster 1 and 3) are stabilized by RNA base stacking, local hydrogen bonding, and a reduced exposure of A_−1_ to the surrounding solvent. In contrast, solvent-exposed states (Cluster 2) create opportunities for direct hydration of the bulged nucleotide and enhanced association of Na^+^ ions with the phosphate backbone and exposed base atoms. The relative stability of the metastable states therefore depends on the balance between burial within the duplex and stabilization by the solvent-ion environment.

This coupling between RNA structure and solvation also has implications for the interpretation of small-molecule splice-site repair. Structural studies have shown that compounds such as SMN-C5 bind near the exon-intron junction, stabilize the bulged adenine, and reinforce the weak SMN2 5’ SS/U1 snRNA duplex [13, 18]. Our simulations suggest that stacked states are already present within the intrinsic ensemble of the unliganded RNA. The estimated penalty for reaching such states is on the range of up to approximately 2 kcal/mol, depending on the solvent model. This energy range is small enough to be plausibly modulated by ligand binding. Therefore, splice-modifying ligands may act by conformational selection rather than by induced-fit. This provides a mechanistic basis for understanding why small molecules that contact the bulged region can have a pronounced effect on SMN2 exon 7 inclusion.

While solvation was expected to influence the equilibrium of the bulged residue, the magnitude of this dependence was unexpected. Although all explicit-solvent models sampled the same conformational families, their relative state populations differed substantially. OPC, TIP4P-Ew, and SPC/E favored stacked conformations as the dominant basin, whereas TIP3P shifted the equilibrium toward solvent-exposed ones. This solvent-exposed state is biologically relevant because inefficient recognition of the SMN2 exon 7 5’ SS has been linked to the presence of the A_−1_ bulge and the resulting discontinuity in the pre-mRNA/U1 snRNA duplex [13, 15]. Thus, TIP3P preferentially stabilizes a state that is structurally consistent with expected defective splice-site geometry of the biological complex. This underscores the importance of the water model selection in simulation of RNA-defect systems.

This interpretation is consistent with molecular simulation studies showing that water molecules around RNA are more than a passive background. Kührová and co-workers showed that different water models can alter RNA hydration patterns and affect canonical A-RNA helical parameters, particularly through changes in water-mediated interactions around the minor groove and 2’-hydroxyl groups [26]. Other work has emphasized that RNA stability depends on the distribution and residence of counterions near the RNA surface, as well as on the structure of the hydration shell [27]. More recent studies further suggest that solvent-model effects can be especially important in structured or flexible RNA systems, where tertiary contacts, local hydration, and ion organization are tightly coupled [28, 29]. Our results extend these observations to a local splice-site defect, showing that even a single bulged nucleotide can create a solvation-sensitive region capable of reshaping the conformational equilibrium.

Several limitations should be considered when interpreting these results. First, although Hamiltonian replica-exchange simulations improve sampling, the conformational landscape of RNA defects may still contain slow, or entropy-driven transitions that are difficult to converge fully. Second, the simulations focus on the RNA duplex environment and do not include the complete spliceosomal context. Third, the observed solvent-model dependence indicates that quantitative state populations should be interpreted cautiously, not only in this system but also in future simulations of RNA defects. Finally, the present analysis describes the unliganded landscape of the defect; direct simulations of ligand-bound systems would be required to determine how individual splicing modifiers redistribute the conformational ensemble.

## Conclusion

In this work, we characterized the intrinsic conformational landscape of the SMN2 A_−1_ splice-site defect, which populates three metastable conformational states that differ in local duplex continuity, stacking interactions, hydrogen-bonding patterns, solvent exposure, and ion association. The recovery of these states across multiple reduced representations supports the view that the SMN2 A_−1_ defect forms a structured conformational ensemble rather than a continuously disordered bulge.

A central finding of this study is that the relative populations of these states are strongly dependent on the explicit-solvent model. Although the same broad conformational families are sampled across water models, their equilibrium weights differ substantially, indicating that hydration and counterion organization are integral components of the RNA defect landscape. Water molecules and Na^+^ ions stabilize solvent-exposed or partially disrupted configurations, whereas RNA-intrinsic stacking and hydrogen-bonding interactions favor more duplex-like states. Thus, the conformational equilibrium of the A_−1_ bulge emerges from a balance between local RNA topology and the surrounding solvent-ion environment.

These results provide a dynamic framework for interpreting small-molecule repair of the weak SMN2 5’ SS. Stacked conformations are already accessible within the intrinsic ensemble of the unliganded RNA, although their populations depend on the solvent representation. Splice-modifying ligands may therefore act by shifting this pre-existing equilibrium toward splice-site-compatible conformations, while also reorganizing hydration and ion contacts around the A_−1_ defect. More broadly, our findings highlight RNA connectivity defects as solvent-sensitive, emphasizing the need to develop better solvation models when modeling or designing RNA-directed therapeutics.

The four-point OPC water model has emerged as a superior choice for molecular dynamics simulations of nucleic acids, largely due to its improved description of water’s liquid-state properties and electrostatic distribution [30]. When integrated with optimized RNA phosphate parameters and the AMBER OL15 force field [31–33], OPC has been shown to significantly improve the stability of secondary structures and the accuracy of conformational sampling compared to traditional three-point models [34]. This combination effectively balances the competing forces of base stacking and backbone solvation, providing a more reliable framework for capturing the subtle structural transitions-such as those observed in the SMN-C5 modifier complex-that are often misrepresented by less sophisticated solvent descriptions.

## Methods

### Molecular Dynamics Simulations

All-atom MD simulations were conducted using the GROMACS 2021.4 software package [35]. The initial nucleic acid coordinates were obtained from the solution NMR structure of the RNA duplex (PDB ID: 6HMO) [13]. The system was solvated in a truncated octahedral box, ensuring a minimum distance of 12 Å between the solute and the box boundaries, and modeled using the AMBER OL15 force field [31–33] To evaluate the impact of the solvent environment, four explicit water models were employed in independent simulation sets: OPC [30], TIP3P [36, 37], SPC/E [38, 39], and TIP4P-Ew [40]. Each system was neutralized and the ionic strength was adjusted to 150 mM NaCl using their respective Joung-Cheatham parameters [41].

To optimize computational efficiency, we utilized a 4 fs time-step for the OPC, TIP4P-Ew, and TIP3P models, and a 3 fs time-step for the SPC/E model, enabled by the use of the hydrogen mass repartitioning [42]. The system preparation involved successive stages of energy minimization and equilibration. Isothermal-Isobaric conditions were maintained using the velocity-rescaling (V-rescale) thermostat [43] and the C-rescale barostat; both with a coupling constant of 10 steps. All bond lengths were constrained using the LINCS algorithm [44]. A 10 Å cutoff was applied to van der Waals and short-range electrostatic interactions, with long-range electrostatics calculated via the Particle Mesh Ewald (PME) method [45].

Hamiltonian replica exchange molecular dynamics (HREMD) was used with PLUMED 2.7.2 [46] to enhance conformational sampling and facilitate the crossing of high-energy barriers [47–49]. For each solvent model, systems were simulated for an aggregated simulation time of 3 *μ*s (Supplementary Figure S17). Systems were set up with 6 replicas, with an inverse temperature scaling corresponding to the following temperatures: 300.00, 320.88, 342.96, 366.44, 391.47, 418.1. The range of temperatures was chosen for promoting the conformational sampling without increasing significantly unfolding, in line to previous works [50]. Exchange attempts were performed every 2500 steps, with successful average exchange rates ranging around 25%.

### Feature Selection

To characterize the conformational landscape of the A_−1_ defect and its surrounding residues, we defined a 22-dimensional feature space comprising geometric and structural descriptors. Hydrogen-bonding stability and alternative base-pairing configurations were monitored via inter-base distances between C_9_-A_−1_, U_10_-G_−1_, and C_9_-G_−1_. To resolve the rotational orientation of these bases, the torsion angles for the C_9_-A_−1_ and U_10_-G_−1_ pairs were decomposed into their sine and cosine components. Base-stacking geometry was quantified for neighboring residues (C_8_-C_9_, C_9_-U_10_, G_−1_-A_−1_, and A_−1_-G_1_) by calculating the angle between the normals of their respective base planes. Furthermore, the SASA of C_9_ and A_−1_ was calculated to assess solvent exposure and local flexibility. All structural features were extracted using the `MDAnalysis` [51, 52] and `MDtraj` [53] Python libraries.

### Dimensionality Reduction and Clustering

To map the dominant conformational states across a unified landscape, the 22-dimensional feature sets extracted from all four water model trajectories were concatenated into a single shared space. Given that the resulting conformational space is high-dimensional, we utilized a comparative framework leveraging both linear and non-linear representation learning methods to isolate the underlying metastable states.

First, standard linear dimensionality reduction was performed using Principal Component Analysis (PCA) [21, 54, 55]. In parallel, multiple deep non-linear encoder-decoder neural network architectures were trained to evaluate their latent spaces as alternative, low-dimensional representations.

To explore the dimensionality-reduction capabilities and topological limits of these neural frameworks, the encoder network dimensions were systematically varied over *d ∈* {2, 4, 6, 8} . This architectural evaluation encompassed deterministic models, Autoencoders (AE)[56], Denoising Autoencoders (DAE)[57], and Contractive Autoencoders (CAE)[58], alongside probabilistic frameworks, including Variational Autoen-coders (VAE)[59], *β*-VAEs (BVAE)[60], Variational Ladder Autoencoders (VLAE)[61], and Denoising VAEs (DVAE)[62]. All models were implemented using the `PyTorch` framework. Detailed hyperparameter optimization protocols and final model selections for these networks are provided in the Supporting Information (Supplementary Methods and Supplementary Tables S1-S4). Hyperparameter optimization was conducted via the `Optuna` framework [63].

Metastable conformational states within each reduced-dimensional representation were partitioned using the HDBSCAN algorithm [22, 64, 65]. Linear dimensionality reduction via PCA, HDBSCAN clustering and t-distributed Stochastic Neighbor Embedding (t-SNE) were performed using the `scikit-learn` library [66]. To visualize the resulting high-dimensional space, t-SNE [67] was applied to compress the coordinates into a clear two-dimensional projection.

### Free Energy Analysis

The relative free energy (Δ*G*_*i*_) of each structural cluster was calculated from its equilibrium population distribution. The discrete probability density of a given cluster, *i*, was mapped onto a free energy using Boltzmann inversion:

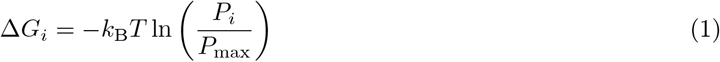

where *k*_B_ is the Boltzmann constant, *T* is the simulation temperature (300 K), *P*_*i*_ represents the equilibrium probability of cluster *i*, and *P*_max_ denotes the probability of the most populated clusters, which correspond as a reference state (Δ*G* = 0).

### Analysis

Base-pairing interactions were classified according to the Leontis-Westhof nomenclature [68], which categorizes hydrogen bonding based on the three distinct interacting edges of RNA bases: Watson-Crick (W), Hoogsteen (H), and Sugar (S). In addition, the classification further accounts for the relative orientation of the glycosidic bonds, distinguishing between cis (c) and trans (t) geometries. Consecutive base-pairs stacking classified according to the MC-annotate classification [69] as upward (3’-5’), downward (5’-3’), outward (5’-5’), and inward (3’-3’). The classification and labeling of the base-pairs interactions and consecutive base-pairs stacking interactions were performed using the `Barnaba` software package [70].

The Hydrogen bonds are defined by a donor–acceptor distance cutoff of 3.5 Åand a minimum angle of 150^°^. All calculations were performed using the `MDAnalysis` library [51, 52]. The RDFs were performed using the `MDAnalysis` Python library [51, 52]. The three-dimensional density maps of sodium ions (*Na*^+^) and water were calculated using a 4 Åcutoff distance using the `MDAnalysis` Python library [51, 52] with a box dimension of 50 Åand a grid spacing of 1.0 Åper cell. The density maps were visualized with Visual Molecular Dynamics (VMD) [71].

## Data and Software Availability

Molecular structures of the three metastable states in PDB format, as well as Jupyter Notebooks with the code used for this work, can be found in a GitHub repository: https://github.com/palominohernandez/SMN2-2026

## Supporting Information

The following files are available free of charge.

- **Supplementary Methods:** Detailed description of the hyperparameter optimization procedure using Optuna, including definitions of the scoring function and evaluation metrics (continuity, trustworthiness, Procrustes similarity, distance correlation, density rank correlation, and validation reconstruction loss). Description of the hyperparameter search space for the dimensionality reduction models.
- **Supplementary Tables:** Table S1: Best model hyperparameters for AE models across latent dimensions. Table S2: Best model hyperparameters for DAE and CAE models. Table S3: Best model hyperparameters for VAE and *β*-VAE models. Table S4: Best model hyperparameters for VLAE and DVAE models. Table S5: Explained variance ratio of the first eight principal components. Table S6: Model-specific hyperparameter search ranges used during optimization. Table S7: Base-pair interactions and probabilities across the three conformational clusters. Table S8: Consecutive base-pair stacking interactions and probabilities across the three conformational clusters.
- **Supplementary Figures:** Figure S1: Feature-principal component correlation matrix for the first eight principal components. Figure S2: Two-dimensional free energy landscape projected onto the first two principal components. Figure S3: Comparison of latent-space representations obtained from VAE, *β*-VAE, VLAE, and DVAE models. Figure S4: Comparison of latent-space representations obtained from PCA, AE, DAE, and CAE models. Figure S5: RMSD and SASA analysis of three RNA conformational clusters. Figure S6: Inclination angle for the three clusters. Figure S7: Probability density distributions of key *O*2^*′*^− *O*2^*′*^ distances across clusters. Figure S8: Three-dimensional water density maps around the RNA for the three clusters and water models. Figure S9: Three-dimensional sodium ion density maps around the RNA for the three clusters and water models. Figure S10: Backbone angle distributions of residues U_10_ and G_11_ in Cluster 3. Figure S11-S13: Radial distribution functions (RDFs) of water oxygen atoms around A_−1_ atoms for Clusters 1-3. Figure S14-S15: Radial distribution functions (RDFs) of sodium ions around A_−1_ atoms for Clusters 1-3. Figure S17: Time-resolved cluster trajectories of replicas. Figure S18: Reconstruction loss for the different model architectures and dimensions.

## Author Contributions

M.K.: Software; Investigation; Formal analysis; Validation; Visualization; Writing – original draft; Writing – review & editing.

L.L.: Software; Investigation; Formal analysis; Visualization; Writing – original draft; Writing – review & editing.

O.P.-H.: Conceptualization; Methodology; Software; Investigation; Formal analysis; Supervision; Funding acquisition; Project administration; Writing – original draft; Writing – review & editing.

## Acknowledgements

This project was funded by the Deutsche Forschungsgemeinschaft (DFG, German Research Foundation – SFB1552 – 465145163). The authors gratefully acknowledge the Gauss Centre for Supercomputing e.V. (www.gauss-centre.eu) for funding this project by providing computing time through the John von Neumann Institute for Computing (NIC) on the GCS Supercomputer JUWELS at Jülich Supercomputing Centre (JSC); application no. 29086. The authors thank Prof. Dr. Paul Czodrowski for his support in this work.

## Notes

M.K. and O.P.-H. contributed equally to this work. The authors declare no competing financial interests.

