## Supplementary material for "Solvation Shapes the Conformational Landscape of a Therapeutically Relevant SMN2 Splice-Site Defect": SI info

### Supporting Information

### Supplementary Methods

#### Loss Functions

Mapping a high-dimensional feature space into a lower-dimensional latent representation can be achieved with various autoencoder (AE) architectures. Each of these AE variations consist of two sequential neural networks: an encoder $f_\theta$, which compresses the input data $x \in \mathbb{R}^D$ into a lower-dimensional latent bottleneck $z \in \mathbb{R}^d$ (where $d < D$), and a decoder $g_\theta$, which attempts to reconstruct the original input from the latent space, producing $\hat{x}$. There are two classes of AE architectures: Deterministic and probabilistic (variational) architectures. Deterministic AEs map each input sample to a fixed point in the latent space, while probabilistic AEs map the input to a continuous probability distribution.

**Autoencoder** The baseline AE is optimized by minimizing the reconstruction loss between input $x$ and its reconstruction $\hat{x}$, calculated here as the Mean Squared Error (MSE) over N samples:

$$\mathcal{L}_{\text{recon}}(x, \hat{x}) = \frac{1}{N} \sum_{i=1}^{N} \left( \frac{1}{D} \sum_{j=1}^{D} (x_{i,j} - \hat{x}_{i,j})^2 \right) \tag{1}$$

**Denoising Autoencoder** A DAE intentionally corrupts the input data by injecting isotropic Gaussian noise $\epsilon \sim \mathcal{N}(0, \sigma^2 I)$ and trains the network to reconstruct the uncorrupted sample. It is trained with the same loss function as a regular AE.

**Contracting Autoencoder** The CAE explicitly regularizes the latent space by penalizing the sensitivity of the encoder's activations to small changes in the input. This is achieved by adding the Frobenius norm of the encoder's Jacobian matrix $J_{f\theta}(x)$ to the loss function:

$$\mathcal{L}_{\text{CAE}} = \mathcal{L}_{\text{recon}}(x, \hat{x}) + \lambda \cdot \mathcal{L}_{\text{contractive}}(x) \tag{2}$$

$$\mathcal{L}_{\text{contractive}}(x) = \frac{1}{N} \sum_{i=1}^{N} \| J_{f\theta}(x_i) \|_F^2 \tag{3}$$

where $\lambda$ is a scaling hyperparameter that balances reconstruction fidelity against the contractive penalty.

**($\beta$-) Variational Autoencoder** Variational Autoencoders (VAEs) are grounded in Bayesian inference and optimize an Evidence Lower Bound (ELBO), which balances the reconstruction loss against a Kullback-Leibler (KL) divergence term. The KL divergence forces the learned latent distribution $Q\mathcal{Q}(z|x)$ to match a predefined prior distribution $\mathcal{P}(z)$. The BVAE adds the hyperparameter $\beta$ to the loss function, which modulates the degree of latent space disentanglement:

$$\mathcal{L}_{\text{VAE}} = \mathcal{L}_{\text{recon}}(x, \hat{x}) + \beta \cdot \mathcal{D}_{\text{KL}}\big(\mathcal{Q}(z|x) \parallel \mathcal{P}(z)\big) \tag{4}$$

**Variational Laplace Autoencoder** The VLAE replaces the standard Gaussian prior with a Laplace distribution. This enforces sparsity in the latent space representations. The KL-divergence of the VLAE is calculated as follows:

$$\mathcal{D}_{\text{KL}}\big(\mathcal{Q}(z|x) \parallel \mathcal{P}(z)\big) = \frac{1}{N} \sum_{i=1}^{N} \sum_{j=1}^{J} \left( \ln\left(\frac{1}{b_{i,j}}\right) + b_{i,j} e^{-\frac{|\mu_{i,j}|}{b_{i,j}}} + |\mu_{i,j}| - 1 \right) \tag{5}$$

where $\mu$ and $b$ are the location and scale parameters of the Laplace posterior.

**Dirichlet Variational Autoencoder** The DVAE assumes a Dirichlet prior, which restricts the latent space to a simplex. The KL divergence for the DVAE is defined using the Gamma ($\Gamma$) and Digamma ($\psi$) functions:

$$\mathcal{D}_{\text{KL}}\big(\mathcal{Q}(z|x) \parallel \mathcal{P}(z)\big) = \sum \log\Gamma(\alpha_i) - \sum \log\Gamma(\hat{\alpha}_i) + \sum(\hat{\alpha}_i - \alpha_i)\psi(\hat{\alpha}_i) \tag{6}$$

where $\alpha_i$are the parameters of the prior distribution and $\hat{\alpha_i}$ are the parameters of the approximate posterior.

#### Hyperparameter Optimization

Hyperparameter optimization (HPO) was done with the Optuna framework [63]. The score that guides the HPO algorithm is defined in equation 8:

$$\text{Score} = 3 \times \text{Continuity} + 3 \times \text{Trustworthiness} + 2 \times \text{Procrustes Similarity} \tag{7}$$
$$+ 2 \times \text{Distance Correlation} + 2 \times \text{Density Matrix Correlation} - 2 \times \text{Validation Reconstruction Loss} \tag{8}$$

**Continuity**

$$\text{Continuity}(k) = 1 - \frac{2}{nk(2n - 3k - 1)} \sum_{i=1}^{n} \sum_{j \in V_k(i)} (\hat{r}(i,j) - k) \tag{9}$$

**Trustworthiness**

$$\text{Trustworthiness}(k) = 1 - \frac{2}{nk(2n - 3k - 1)} \sum_{i=1}^{n} \sum_{j \in U_k(i)} \max(r(i,j) - k) \tag{10}$$

**Procrustes Similarity** Given two centered and normalized embedding matrices $A, B \in \mathbb{R}^{n \times d}$, the orthogonal Procrustes problem seeks an orthogonal rotation matrix $R \in \mathbb{R}^{d \times d}$ that minimizes the squared Frobenius norm of the residual differences:

$$\min_R \|A - BR\|_F^2 \quad \text{subject to} \quad R^T R = I_d \tag{11}$$

Using the Singular Value Decomposition (SVD) of the cross-covariance matrix $B^T A = U \Sigma V^T$, the optimal rotation matrix is solved via:

$$\hat{R} = UV^T \tag{12}$$

The structural discrepancy is measured by the residual Mean Squared Error (disparity):

$$\text{Disparity} = \frac{1}{n}\|A - B\hat{R}\|_F^2 \tag{13}$$

The final Procrustes similarity metric is bounded as:

$$\text{Procrustes Similarity} = \max(0.0, 1.0 - \text{Disparity}) \tag{14}$$

**Distance Correlation** Let $d_1$ and $d_2$ be the flattened vectors containing all $N = \frac{n(n-1)}{2}$ pairwise Euclidean distances calculated within embeddings $Z_1$ and $Z_2$ respectively. The Spearman rank correlation coefficient $\rho$ evaluates global distance preservation by computing:

$$\rho = 1 - \frac{6\sum_{i=1}^{N}\left(\text{rank}(d_{1,i}) - \text{rank}(d_{2,i})\right)^2}{N(N^2 - 1)} \tag{15}$$

**Density Rank Correlation** In a $d$-dimensional latent space, the local log-density proxy $s_i$ for a specific data point $i$ is calculated using the distance to its $k$-th nearest neighbor, denoted as $r_k(i)$:

$$s_i = -d \cdot \log(r_k(i) + \epsilon) \tag{16}$$

where $\epsilon = 10^{-12}$ prevents numerical instability. Given density proxy vectors $s^{(1)}$ and $s^{(2)}$ extracted from two independent seed runs, the stability of the density profile across the manifold is measured using the Spearman rank correlation:

$$\text{Density Rank Corr} = 1 - \frac{6\sum_{i=1}^{n}\left(\text{rank}(s_i^{(1)}) - \text{rank}(s_i^{(2)})\right)^2}{n(n^2-1)} \tag{17}$$

**Validation Reconstruction Loss** The validation reconstruction loss in Equation 1 measures the model's ability to reconstruct a datapoint from its latent representation. The minimized validation losses achieved across the different model architectures and latent dimensions are presented in Figure S18. Deterministic autoencoder architectures (AE, DAE, and CAE) with 8 dimensions consistently reproduced the input latent space more effectively and with lower reconstruction error than the variational models.

The search space consisted of general hyperparameters and model-specific parameters. The general hyperparameters included the learning rate, weight decay, and the layer composition of the encoder network. For all models, the decoder network architecture was a symmetric, mirrored version of its corresponding encoder.

### Supplementary Tables

Table S1: Best model hyperparameters for Autoencoder (AE), determined by hyperparameter optimization with Optuna.

| Model | AE | | |
|---|---|---|---|
| Latent Dim | $HD_{max}$ | LR | WD |
| 2 | 32 | $1.98 \cdot 10^{-3}$ | $2.87 \cdot 10^{-5}$ |
| 4 | 32 | $8.25 \cdot 10^{-4}$ | $8.98 \cdot 10^{-5}$ |
| 6 | 32 | $2.16 \cdot 10^{-3}$ | $4.07 \cdot 10^{-5}$ |
| 8 | 32 | $2.93 \cdot 10^{-3}$ | $6.06 \cdot 10^{-5}$ |

Table S2: Best model hyperparameters for Autoencoders DAE and CAE, determined by hyperparameter optimization with Optuna.

| Model | DAE | | | | CAE | | | |
|---|---|---|---|---|---|---|---|---|
| Latent Dim | $HD_{max}$ | LR | WD | Noise Std. | $HD_{max}$ | LR | WD | $\lambda$ |
| 2 | 32 | $1.77 \cdot 10^{-3}$ | $7.84 \cdot 10^{-5}$ | 0.5 | 32 | $2.98 \cdot 10^{-3}$ | $7.04 \cdot 10^{-5}$ | $6.93 \cdot 10^{-5}$ |
| 4 | 32 | $2.93 \cdot 10^{-3}$ | $3.12 \cdot 10^{-5}$ | 0.03 | 32 | $9.40 \cdot 10^{-4}$ | $1.41 \cdot 10^{-5}$ | $4.49 \cdot 10^{-5}$ |
| 6 | 32 | $2.99 \cdot 10^{-3}$ | $4.39 \cdot 10^{-5}$ | 0.25 | 32 | $2.32 \cdot 10^{-3}$ | $4.88 \cdot 10^{-5}$ | $3.52 \cdot 10^{-6}$ |
| 8 | 32 | $2.99 \cdot 10^{-3}$ | $3.82 \cdot 10^{-5}$ | 0.03 | 32 | $3.00 \cdot 10^{-3}$ | $9.93 \cdot 10^{-5}$ | $3.50 \cdot 10^{-6}$ |

Table S3: Best model hyperarameters for VAE and $\beta$-VAE, determined by hyperparameter optimization with Optuna.

| Model | VAE | | | BVAE | | | |
|---|---|---|---|---|---|---|---|
| Latent Dim | $HD_{max}$ | LR | WD | $HD_{max}$ | LR | WD | $\beta$ |
| 2 | 32 | $2.97 \cdot 10^{-3}$ | $5.17 \cdot 10^{-5}$ | 32 | $2.31 \cdot 10^{-3}$ | $5.42 \cdot 10^{-5}$ | 1 |
| 4 | 16 | $1.00 \cdot 10^{-6}$ | $3.03 \cdot 10^{-5}$ | 32 | $1.00 \cdot 10^{-6}$ | $1.37 \cdot 10^{-5}$ | 100 |
| 6 | 32 | $2.02 \cdot 10^{-3}$ | $5.37 \cdot 10^{-5}$ | 32 | $1.01 \cdot 10^{-6}$ | $7.74 \cdot 10^{-5}$ | 4 |
| 8 | 32 | $1.00 \cdot 10^{-6}$ | $4.44 \cdot 10^{-5}$ | 16 | $1.01 \cdot 10^{-6}$ | $2.34 \cdot 10^{-5}$ | 4 |

Table S4: Best model hyperparameters for VLAE and DVAE, determined by hyperparameter optimization with Optuna.

| Model | VLAE | | | | DVAE | | | | |
|---|---|---|---|---|---|---|---|---|---|
| Latent Dim | $HD_{max}$ | LR | WD | $\beta$ | $HD_{max}$ | LR | WD | $\alpha_\Gamma$ | $\beta_\Gamma$ |
| 2 | 32 | $2.67 \cdot 10^{-3}$ | $8.33 \cdot 10^{-5}$ | 1 | 32 | $1.01 \cdot 10^{-6}$ | $3.69 \cdot 10^{-5}$ | 8 | 4 |
| 4 | 32 | $1.51 \cdot 10^{-3}$ | $1.43 \cdot 10^{-5}$ | 1 | 16 | $1.00 \cdot 10^{-6}$ | $9.87 \cdot 10^{-5}$ | 2 | 4 |
| 6 | 16 | $1.00 \cdot 10^{-6}$ | $2.50 \cdot 10^{-5}$ | 50 | 16 | $1.00 \cdot 10^{-6}$ | $3.97 \cdot 10^{-5}$ | 8 | 4 |
| 8 | 16 | $1.00 \cdot 10^{-6}$ | $2.52 \cdot 10^{-5}$ | 8 | 16 | $1.00 \cdot 10^{-6}$ | $2.26 \cdot 10^{-5}$ | 4 | 2 |

Table S5: Explained variance ratio for the first 8 principal components.

| **Component** | **Explained Variance Ratio (%)** |
|---|---|
| PC1 | 32.81 |
| PC2 | 14.39 |
| PC3 | 10.90 |
| PC4 | 6.15 |
| PC5 | 5.27 |
| PC6 | 5.04 |
| PC7 | 4.29 |
| PC8 | 4.13 |
| **Total (8 components)** | **82.98** |

Table S6: Model-specific parameters to optimize.

| AE | DAE | CAE | VAE | BVAE | VLAE | DVAE | |
|---|---|---|---|---|---|---|---|
| - | Noise | $\lambda$ | - | $\beta$ | $\beta$ | $\alpha$ | $\beta$ |
| - | 0.01, 0.03, 0.05, 0.08, 0.10, 0.12, 0.25, 0.4, 0.5 | $10^{-6} - 10^{-1}$ | - | 0.1, 0.25, 0.5, 0.75 | 1.0, 2.0, 4.0, 8.0, 25, 50, 75, 100 | 1.0, 2.0, 4.0, 8.0 | 1.0, 2.0, 4.0, 8.0 |

Table S7: Base-pair interactions and their probabilities across the three clusters. Residue 1 (Res1) belongs to the U1 snRNA, and Residue 2 (Res2) belongs to the SMN2 5' SS. Base pairs are classified according to the Leontis-Westhof scheme: W, H, and S denote Watson-Crick, Hoogsteen, and Sugar edges, respectively, while c/t indicate cis or trans orientation.

| Base-Pair | | Cluster 1 | | Cluster 2 | | Cluster 3 | |
|---|---|---|---|---|---|---|---|
| Res1 | Res2 | Classification | Prob | Classification | Prob | Classification | Prob |
| $A_1$ | $U_8$ | WWc | 0.893 | WWc | 0.867 | WWc | 0.849 |
| $U_2$ | $C_7$ | WWc | 0.991 | WWc | 0.980 | WWc | 0.988 |
| $A_3$ | $U_6$ | WWc | 0.945 | WWc | 0.937 | WWc | 0.940 |
| $C_4$ | $G_5$ | WWc | 0.995 | WWc | 0.998 | WWc | 0.993 |
| $U_5$ | $A_4$ | WWc | 0.996 | WWc | 0.997 | WWc | 0.997 |
| $U_6$ | $A_3$ | WWc | 0.998 | WWc | 0.997 | WWc | 0.997 |
| $A_7$ | $U_2$ | WWc | 0.991 | WWc | 0.996 | WWc | 0.997 |
| $C_8$ | $G_1$ | WWc | 0.978 | WWc | 1.000 | WWc | 1.000 |
| $C_9$ | $A_{-1}$ | – | – | SHc | 0.733 | WWc | 0.909 |
| $C_9$ | $G_{-2}$ | WWc | 1.000 | WWc | 1.000 | – | – |
| $U_{10}$ | $G_{-2}$ | – | – | – | – | WWc | 0.925 |
| $U_{10}$ | $G_{-3}$ | WWc | 0.941 | WWc | 0.985 | – | – |

Table S8: Consecutive Base-pair stacking interactions and their probabilities across the three clusters. The upper section profiles the stacking steps along the U1 snRNA, while the lower section details consecutive steps along the SMN2 5'SS. Stacking alignments are classified as upward (3'–5'), downward (5'–3'), outward (5'–5'), and inward (3'–3').

| Base-Pair | | Cluster 1 | | Cluster 2 | | Cluster 3 | |
|---|---|---|---|---|---|---|---|
| Res1 | Res2 | Classification | Prob | Classification | Prob | Classification | Prob |
| $A_1$ | $U_2$ | Upward | 0.972 | Upward | 0.930 | Upward | 0.923 |
| $U_2$ | $A_3$ | Upward | 0.401 | Upward | 0.337 | Upward | 0.394 |
| $A_3$ | $C_4$ | Upward | 0.970 | Upward | 0.961 | Upward | 0.968 |
| $C_4$ | $U_5$ | Upward | 0.808 | Upward | 0.762 | Upward | 0.761 |
| $U_5$ | $U_6$ | Upward | 0.841 | Upward | 0.861 | Upward | 0.854 |
| $A_7$ | $C_8$ | Upward | 0.995 | Upward | 0.999 | Upward | 0.999 |
| $C_8$ | $C_9$ | Upward | 0.463 | Upward | 0.996 | Upward | 0.494 |
| $C_9$ | $U_{10}$ | Upward | 0.993 | Upward | 0.985 | Upward | 0.884 |
| $U_{10}$ | $G_{11}$ | – | – | – | – | Upward | 0.354 |
| $G_{-3}$ | $G_{-2}$ | Upward | 0.447 | Upward | 0.380 | Upward | 0.455 |
| $G_{-2}$ | $A_{-1}$ | Upward | 0.329 | – | – | Upward | 0.675 |
| $A_{-1}$ | $G_1$ | Upward | 0.950 | – | – | – | – |
| $G_1$ | $U_2$ | Upward | 1.000 | Upward | 1.000 | Upward | 0.999 |
| $A_3$ | $A_4$ | Upward | 0.850 | Upward | 0.731 | Upward | 0.750 |
| $A_4$ | $G_5$ | Upward | 0.652 | Upward | 0.684 | Upward | 0.632 |
| $G_5$ | $U_6$ | Upward | 0.997 | Upward | 0.996 | Upward | 0.996 |
| $U_6$ | $C_7$ | Upward | 0.619 | Upward | 0.635 | Upward | 0.616 |
| $C_7$ | $U_8$ | Upward | 0.726 | Upward | 0.736 | Upward | 0.712 |

#### Supplementary Figures

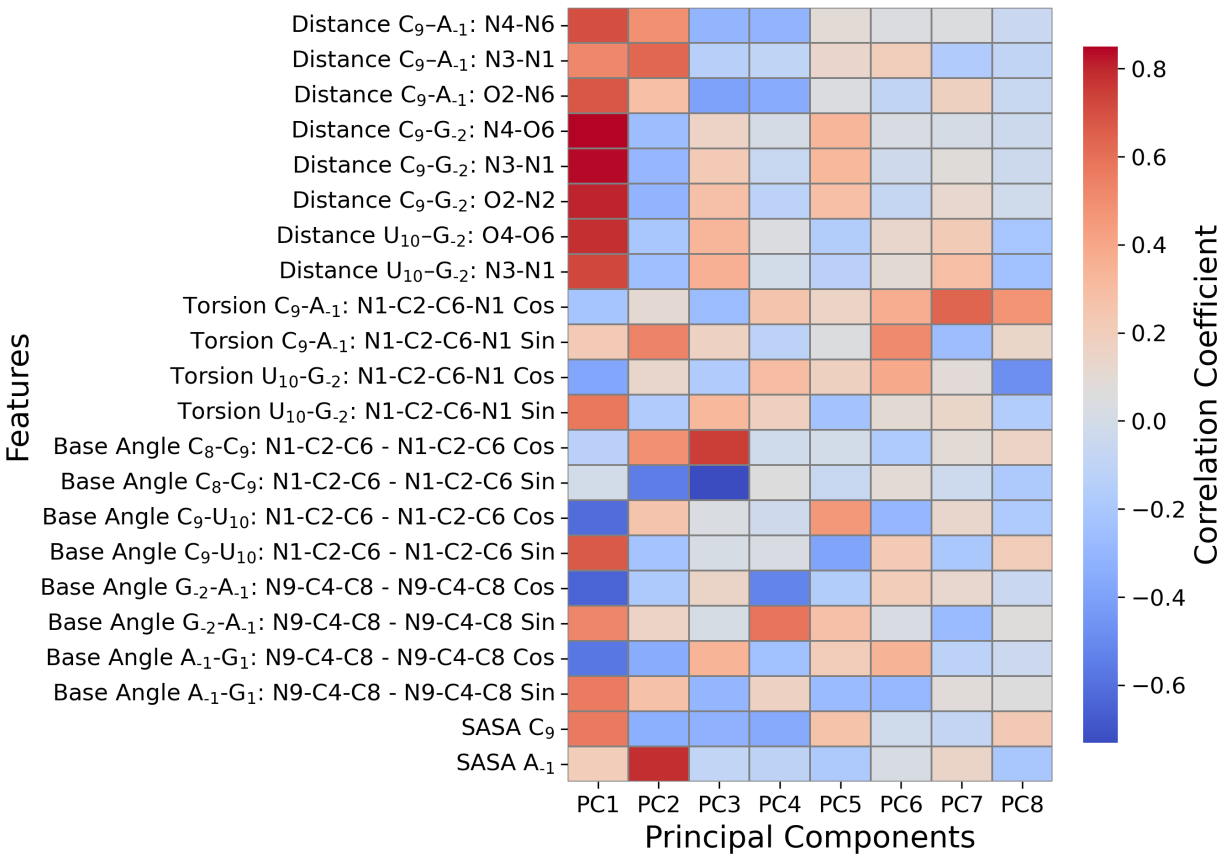


Figure S1: Feature-PC correlations for the first 8 principal components, shown as a matrix. The color bar indicates the correlation coefficient.

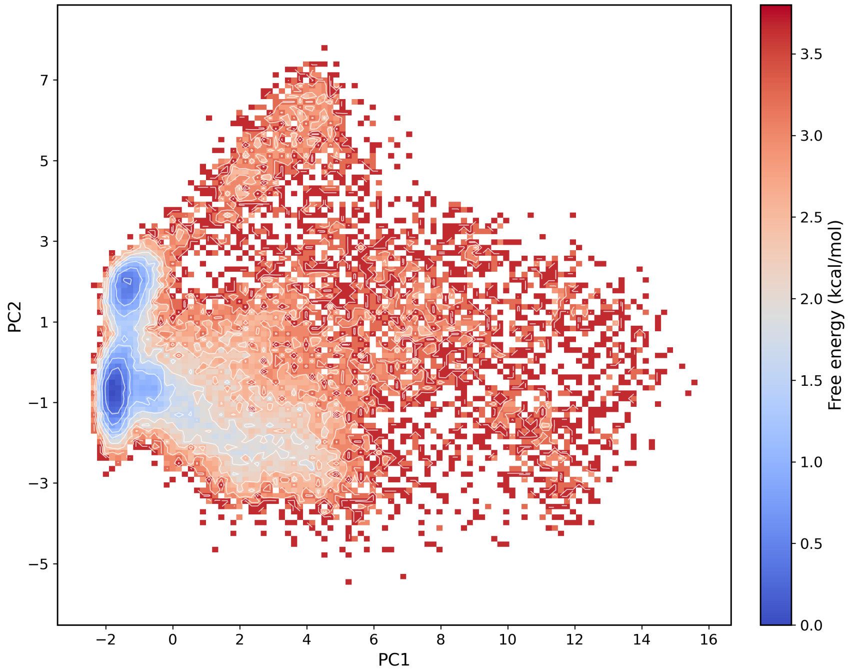


Figure S2: 2D Free Energy Landscape (FEL) of the SMN2 $A_{-1}$ bulge from the combined dataset for OPC, TIP4P-Ew, TIP3P, and SPC/E water models, projected onto the first two principal components (PC1 and PC2). The color bar indicates free energy in kcal/mol.

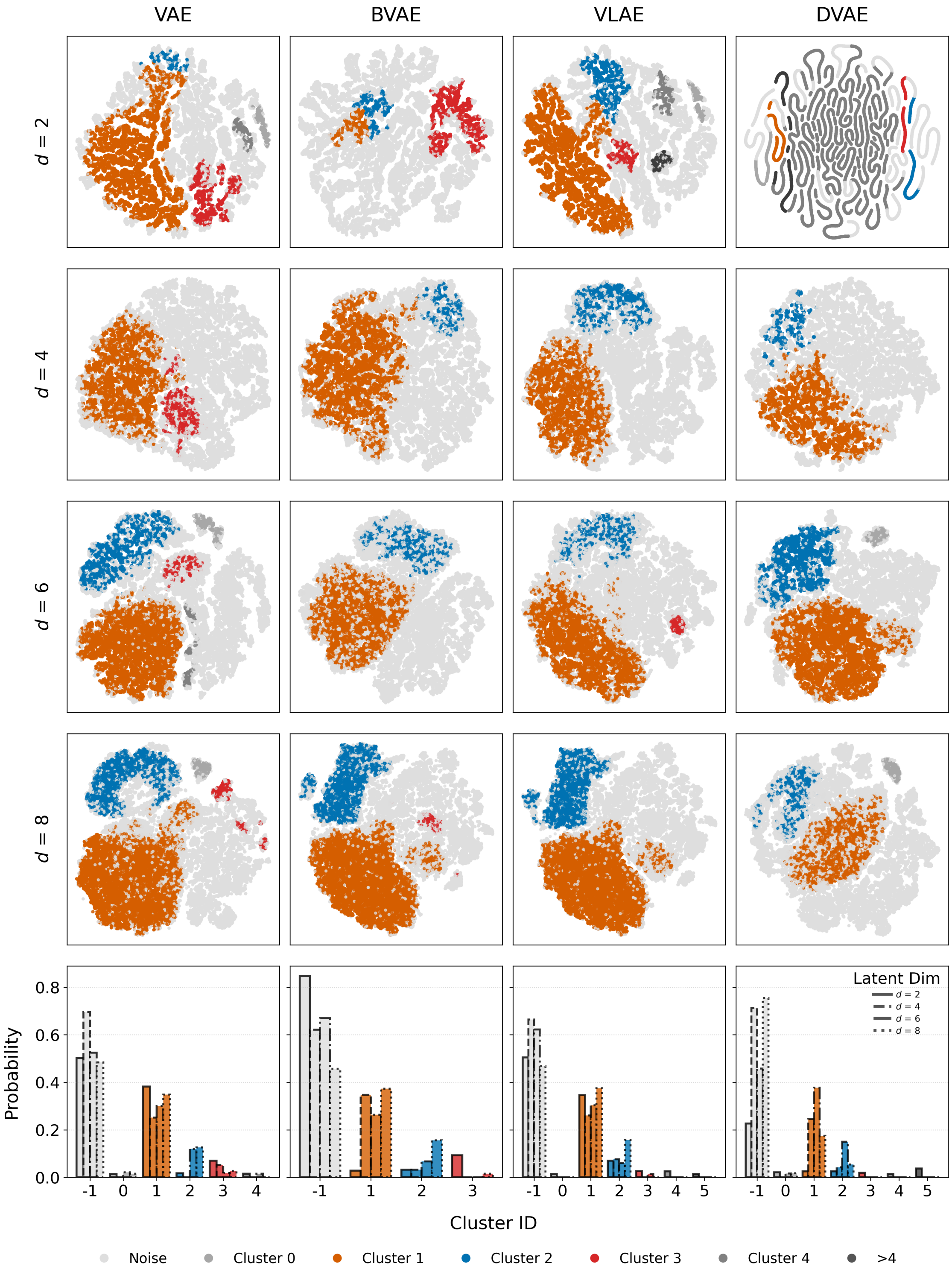


Figure S3: Two-dimensional projections of the conformational space generated via $t$-SNE applied to the latent spaces ($d = 2, 4, 6, 8$) of VAE, BVAE, VLAE, and DVAE. Cluster probabilities extracted across the corresponding embedding dimensions ($d = 2, 4, 6, 8$). Clusters are color-coded as follows: Cluster 1 (orange), Cluster 2 (blue), and Cluster 3 (red), while noise clusters are shown in shades of gray. Cluster IDs >4 are omitted for clarity.

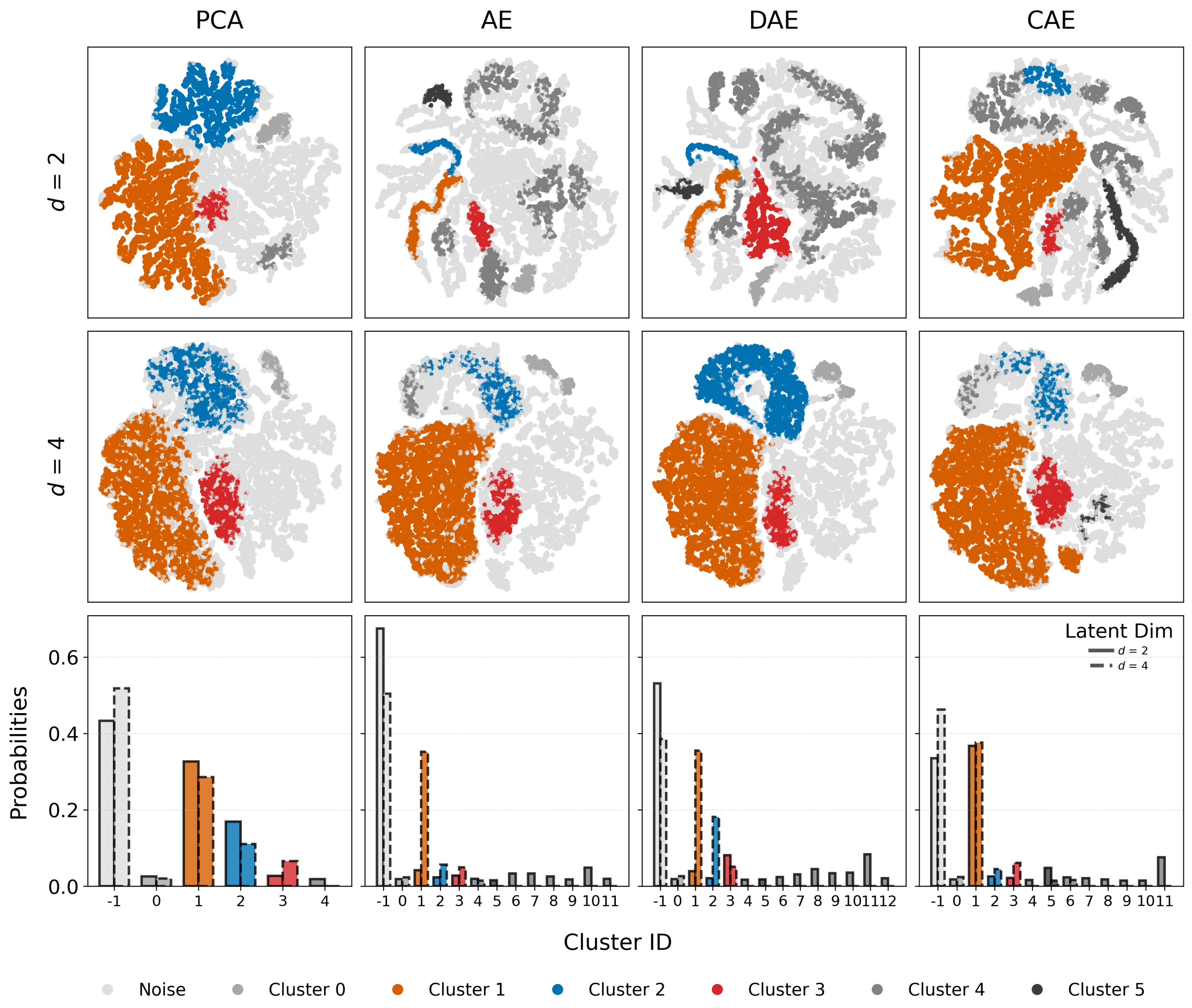


Figure S4: Two-dimensional projections of the conformational space generated via $t$-SNE applied to the latent spaces ($d = 2$ and $d = 4$) of PCA, AE, DAE, and CAE. Cluster probabilities extracted across the corresponding embedding dimensions ($d = 2, 4$). Clusters are color-coded as follows: Cluster 1 (orange), Cluster 2 (blue), and Cluster 3 (red), while noise clusters are shown in shades of gray.

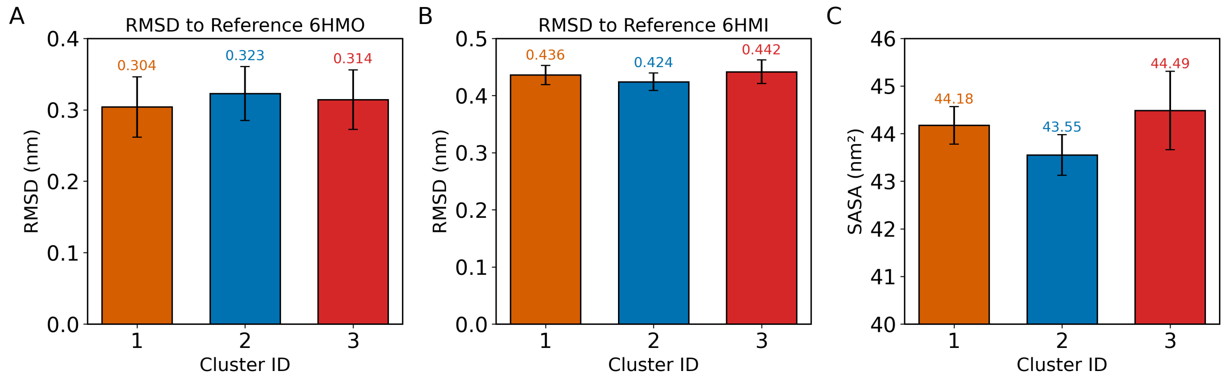


Figure S5: Average RNA heavy-atom root-mean-square deviations (RMSDs) of the three clusters calculated after structural alignment against the reference experimental structures (A) PDB ID: 6HMO and (B) PDB ID: 6HMI. (C) The average solvent-accessible surface area (SASA) of all RNA atoms of the clusters. Error bars represent standard deviations.

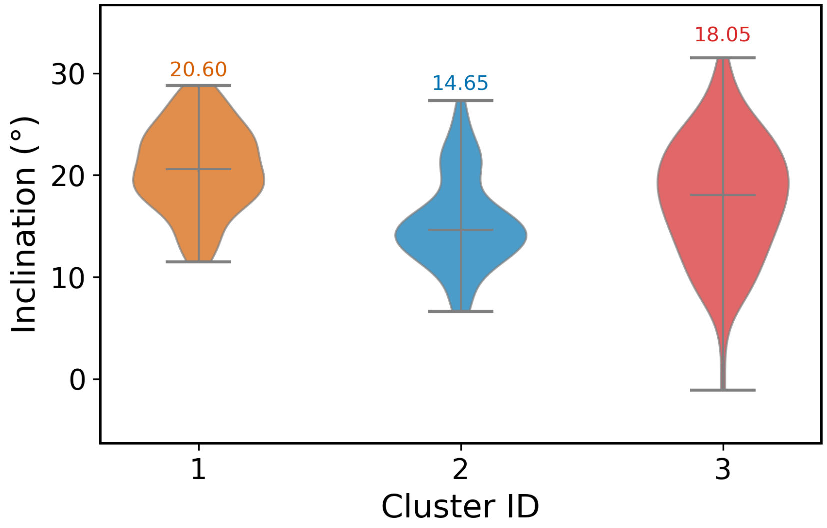


Figure S6: Inclination (rotation around the X-axis) for the three clusters. The text labels indicate the median values. The base inclination was calculated using CURVES+ (version 3.0) [72]. For each cluster, 100 randomly selected frames were analyzed. The RNA duplex was defined as two strands with sequences 5'–3': AUACUUAC-CUG and 3'–5': UCUGAAUGAGG. Residue $A_{-1}$ was treated as unpaired and was therefore excluded from base-pairing analysis.

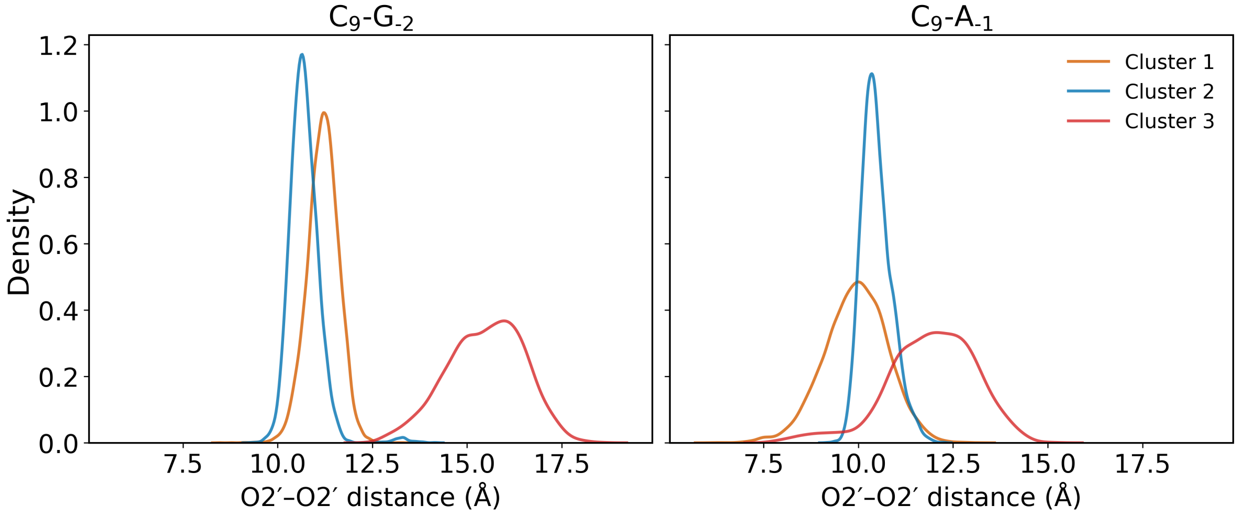


Figure S7: Probability density distributions of $O2' - O2'$ distances for key residue pairs ($C_9$-$G_{-2}$ and $C_9$-$A_{-1}$) across the four clusters. The 2′-hydroxyl group is known to influence RNA conformational dynamics by affecting backbone flexibility.

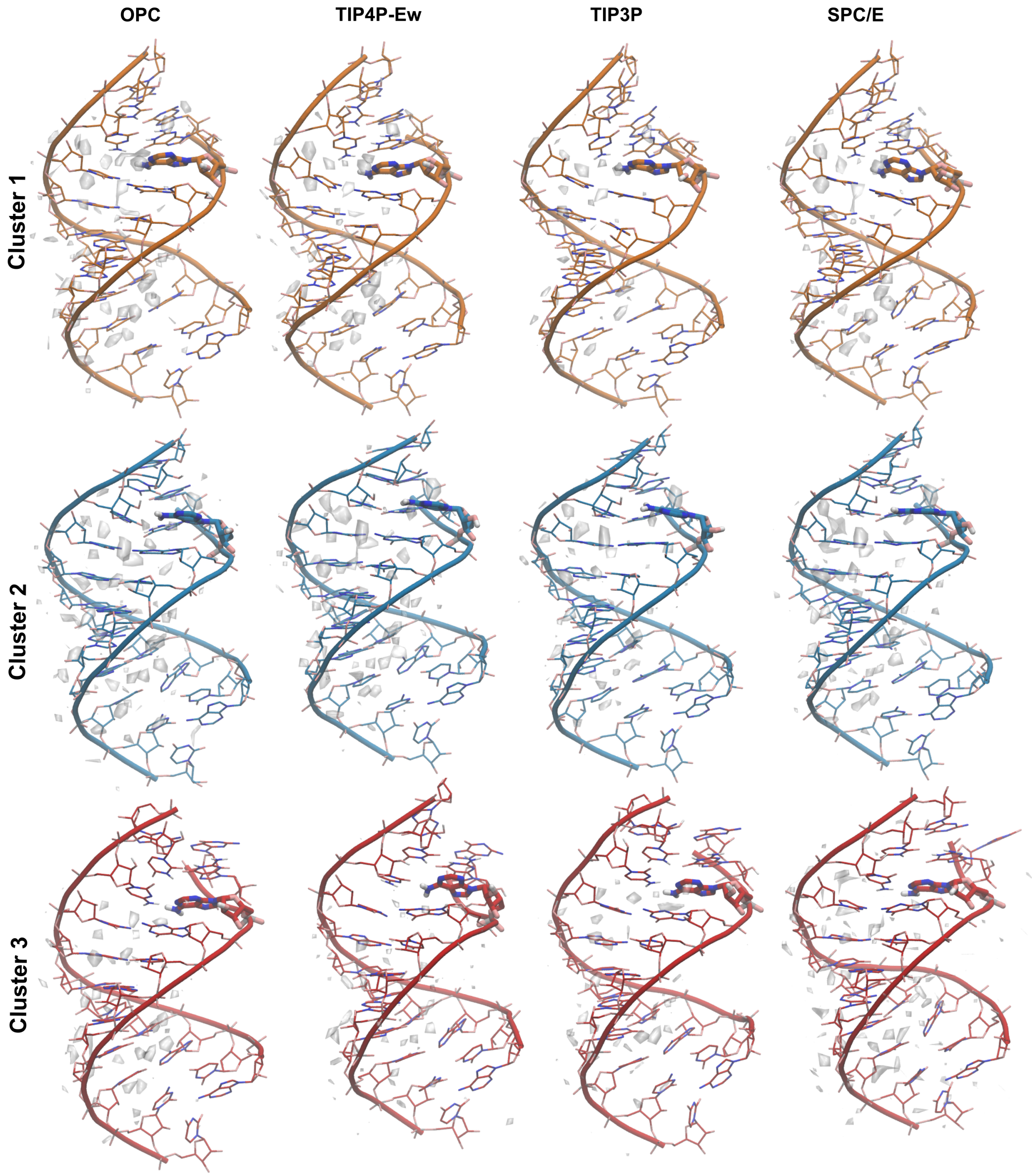


Figure S8: Three-dimensional density maps of water around the RNA were calculated for the three clusters across the four water models (OPC, TIP4P-Ew, TIP3P, and SPC/E). The density maps were visualized using VMD and displayed at the same isocontour level of 0.07 (0.08 for Cluster 3 of TIP4P-Ew and SPC/E models). The water density maps are shown in white.

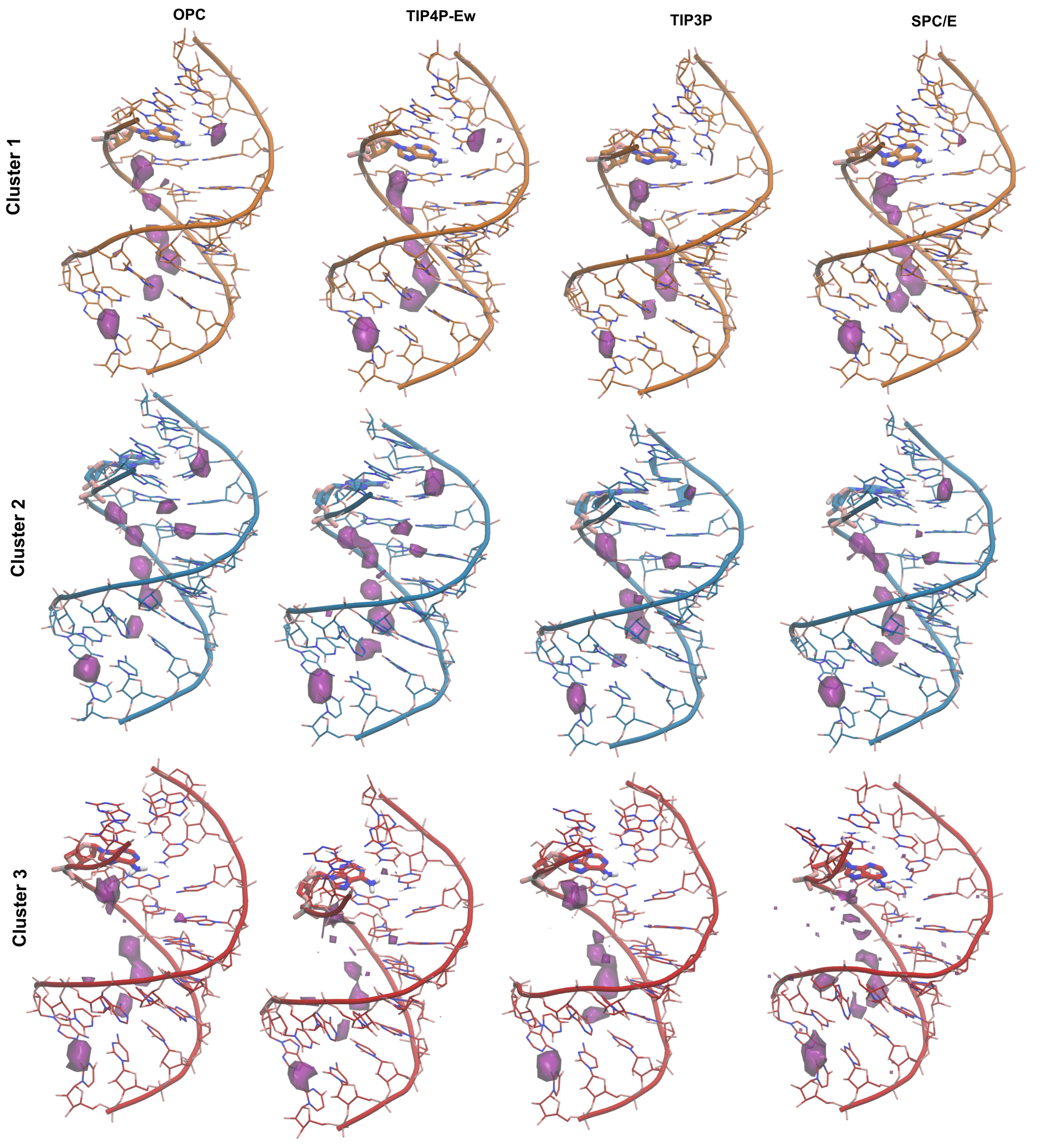


Figure S9: Three-dimensional density maps of sodium ions ($Na^{+}$) around the RNA were calculated for the three clusters across the four water models (OPC, TIP4P-Ew, TIP3P, and SPC/E). The density maps were visualized using VMD and displayed at the same isocontour level of 0.009 (0.018 for Cluster 3 of TIP4P-Ew and SPC/E models). The $Na^{+}$ density maps are shown in purple.

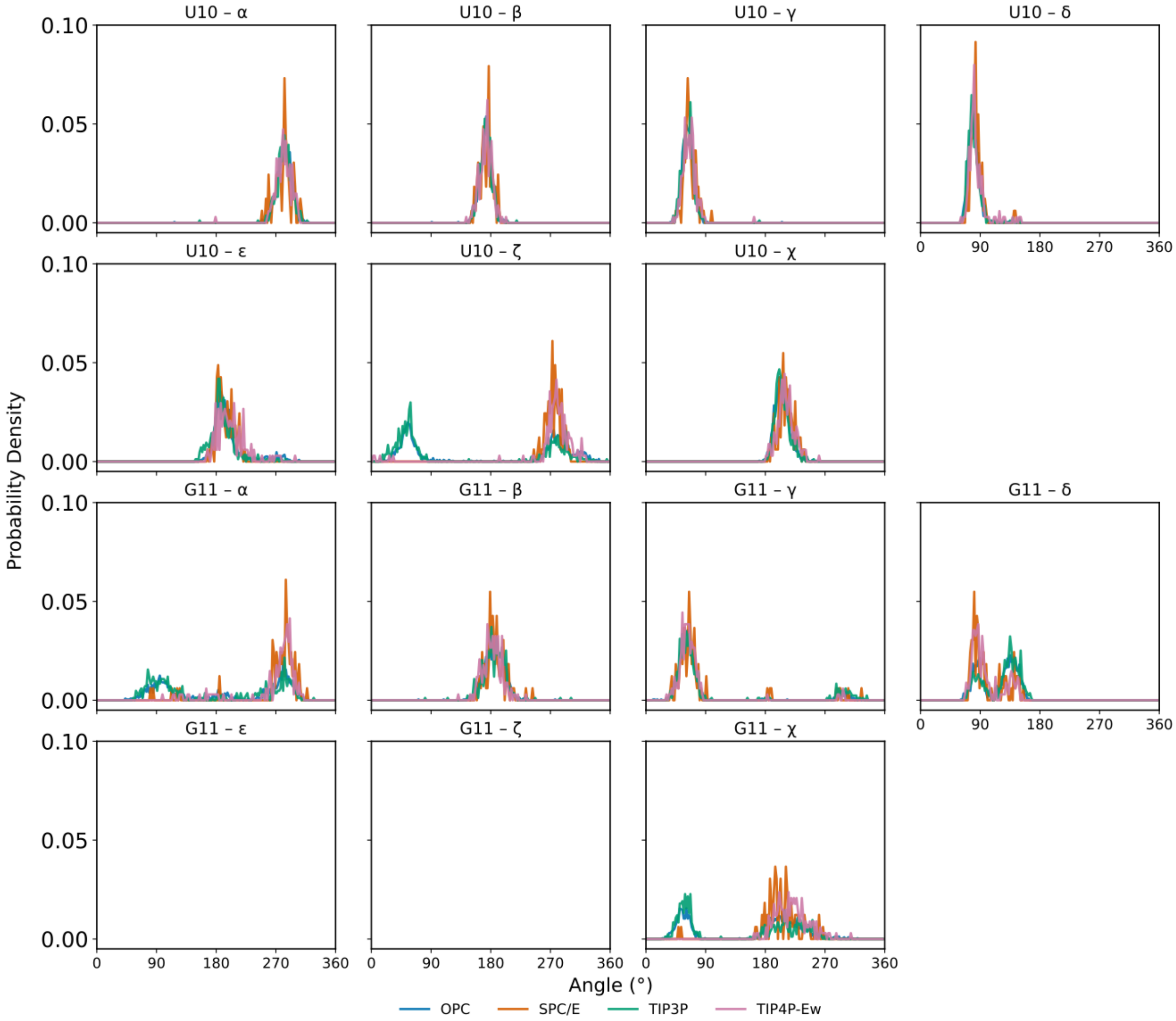


Figure S10: The backbone angle distributions of $U_{10}$ and $G_{11}$ of Cluster 3.

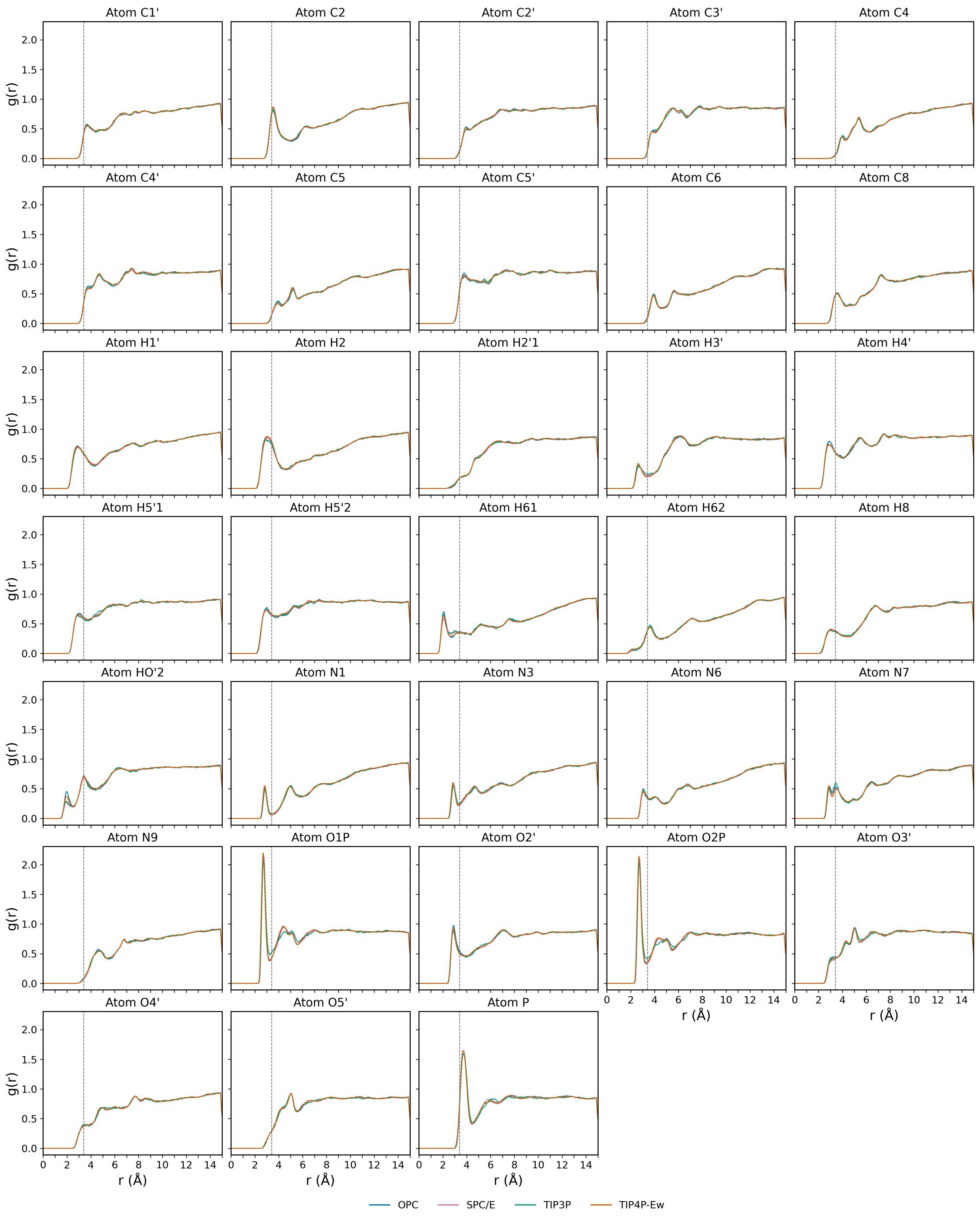


Figure S11: Radial distribution functions (RDFs) of water oxygen ($O_w$) around the $\mathrm{A}_{-1}$ atoms for Cluster 1. The plots show the average RDF, $g(r)$, for all Clusters across the four water models. A dashed black vertical line at 3.4 Åindicates the boundary of the first solvation shell. Water models are distinguished by color: OPC (green), TIP4P-Ew (orange), TIP3P (blue), and SPC/E (purple).

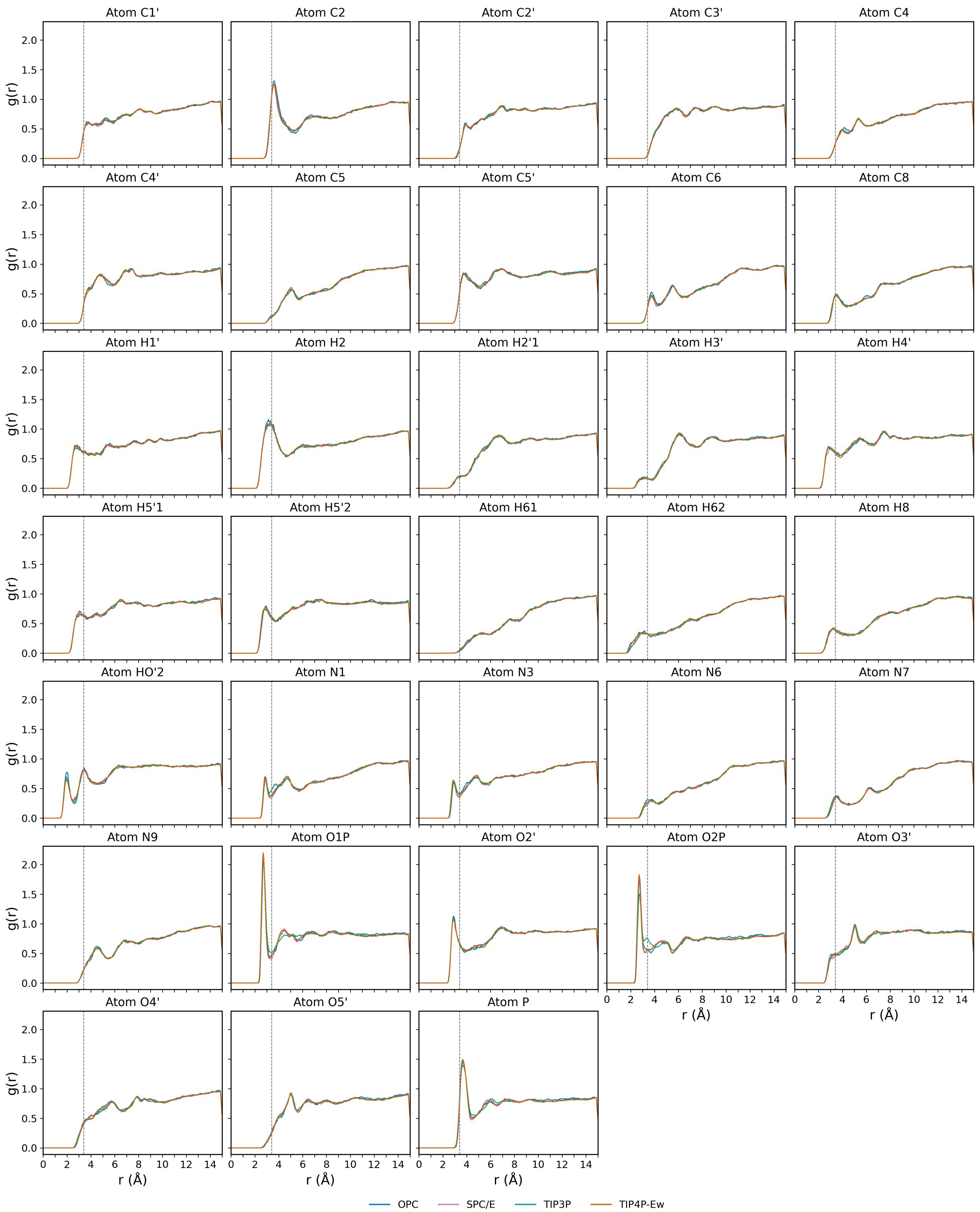


Figure S12: Radial distribution functions (RDFs) of water oxygen ($O_w$) around the $\mathrm{A}_{-1}$ atoms for Cluster 2. The plots show the average RDF, $g(r)$, for all Clusters across the four water models. A dashed black vertical line at 3.4 Åindicates the boundary of the first solvation shell. Water models are distinguished by color: OPC (green), TIP4P-Ew (orange), TIP3P (blue), and SPC/E (purple).

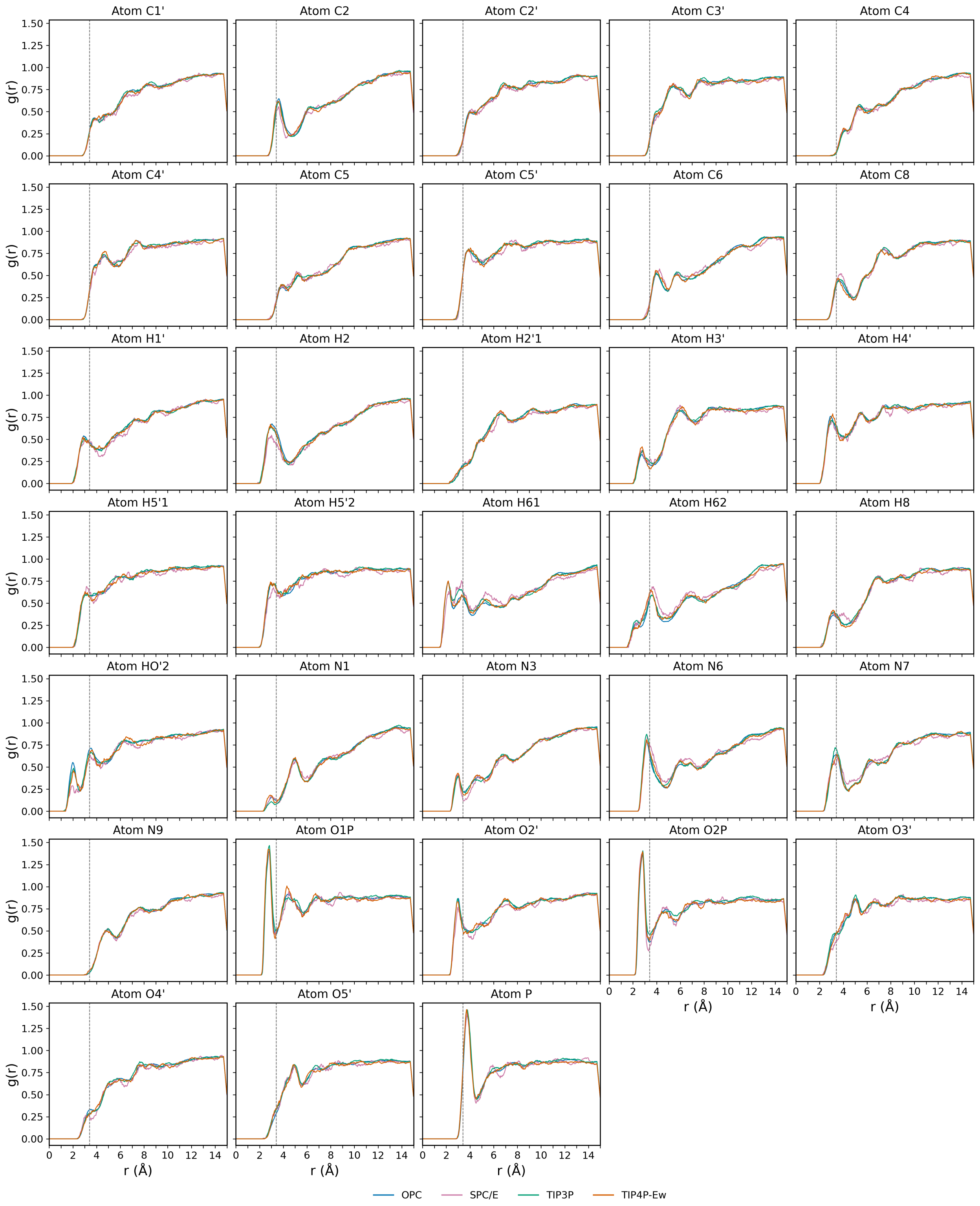


Figure S13: Radial distribution functions (RDFs) of water oxygen ($O_w$) around the $A_{-1}$ atoms for Cluster 3. The plots show the average RDF, $g(r)$, for all Clusters across the four water models. A dashed black vertical line at 3.4 Åindicates the boundary of the first solvation shell. Water models are distinguished by color: OPC (green), TIP4P-Ew (orange), TIP3P (blue), and SPC/E (purple).

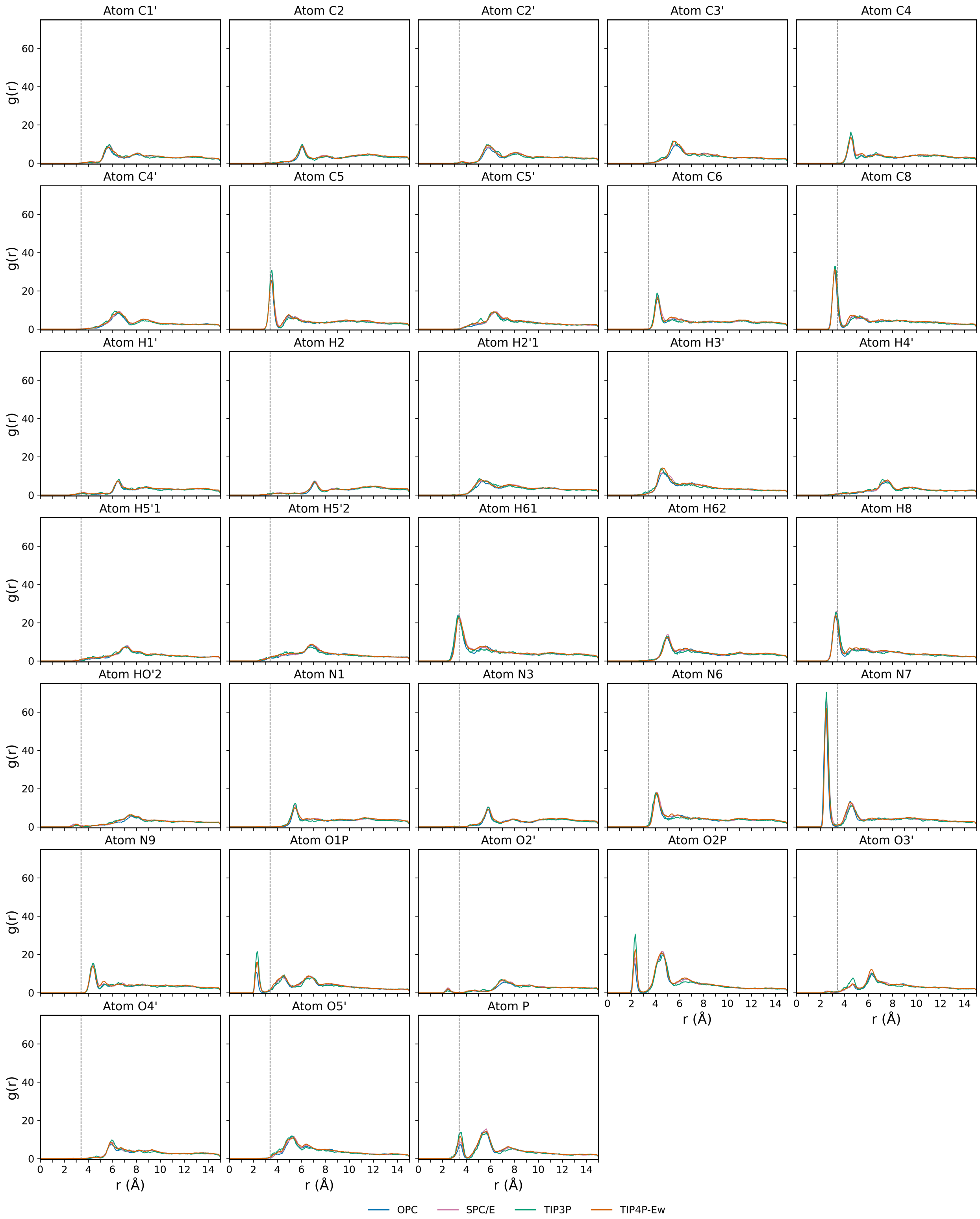


Figure S14: Radial distribution functions (RDFs) of $Na^+$ ions around the $\mathrm{A}_{-1}$ atoms for Cluster 1. The plots show the average RDF, $g(r)$, for all Clusters across the four water models. A dashed black vertical line at 3.4 Åindicates the boundary of the first solvation shell. Water models are distinguished by color: OPC (green), TIP4P-Ew (orange), TIP3P (blue), and SPC/E (purple).

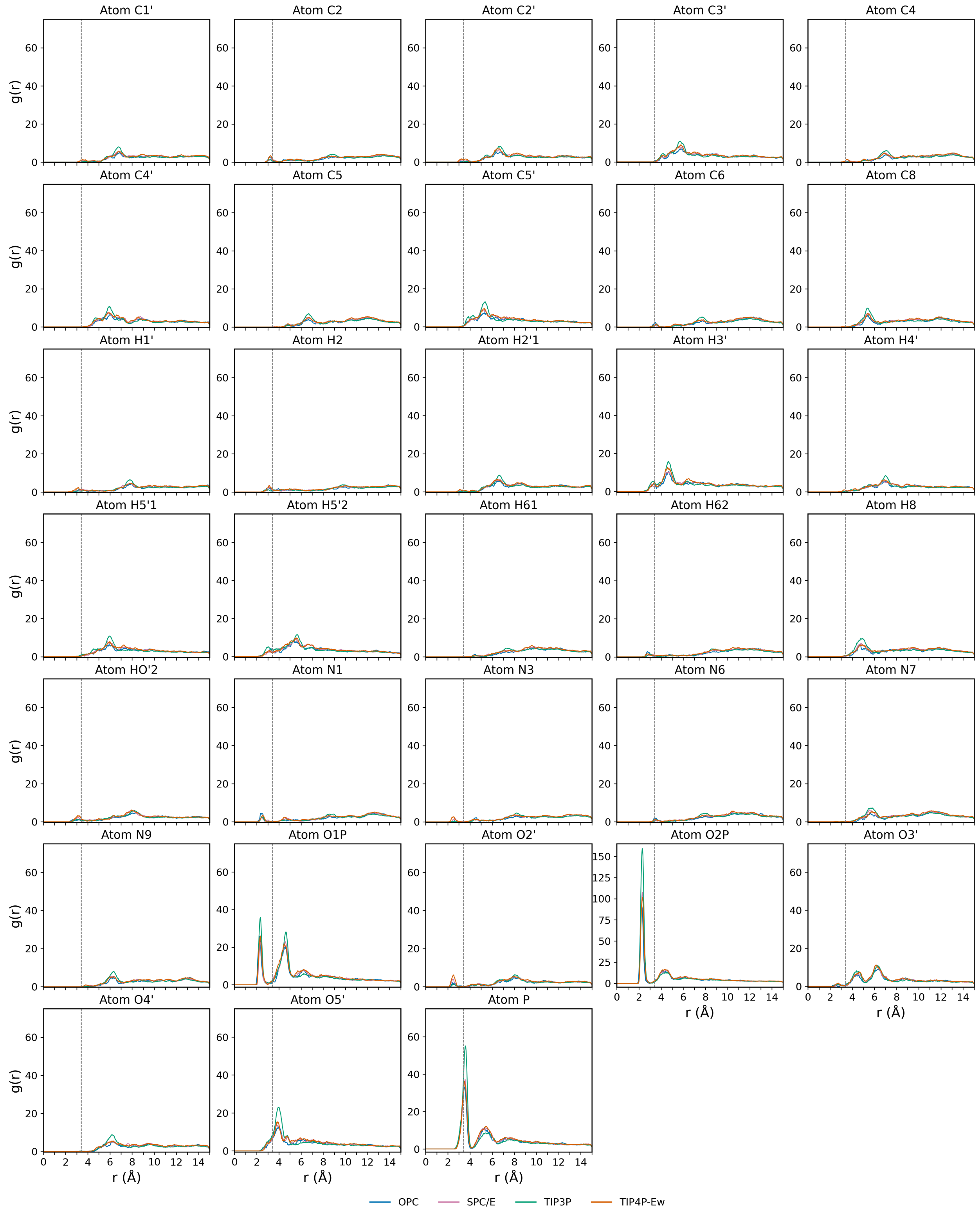


Figure S15: Radial distribution functions (RDFs) of $Na^+$ ions around the $A_{-1}$ atoms for Cluster 2. The plots show the average RDF, $g(r)$, for all Clusters across the four water models. A dashed black vertical line at 3.4 Åindicates the boundary of the first solvation shell. Water models are distinguished by color: OPC (green), TIP4P-Ew (orange), TIP3P (blue), and SPC/E (purple). Note that the y-scale of atom O2P and Cluster 3 is different for clarity.

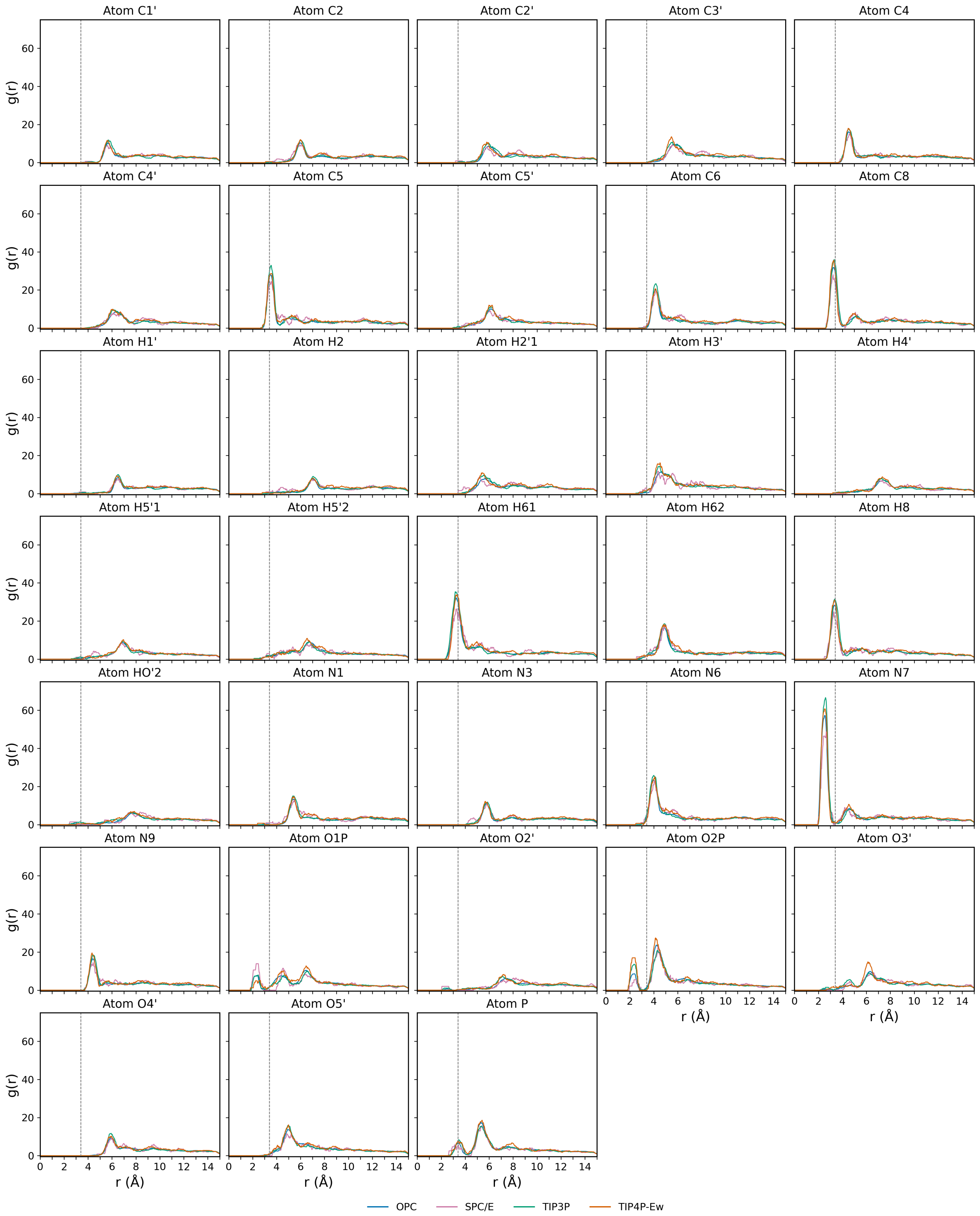


Figure S16: Radial distribution functions (RDFs) of $Na^+$ ions around the $\mathrm{A}_{-1}$ atoms for Cluster 3. The plots show the average RDF, $g(r)$, for all Clusters across the four water models. A dashed black vertical line at 3.4 Åindicates the boundary of the first solvation shell. Water models are distinguished by color: OPC (green), TIP4P-Ew (orange), TIP3P (blue), and SPC/E (purple).

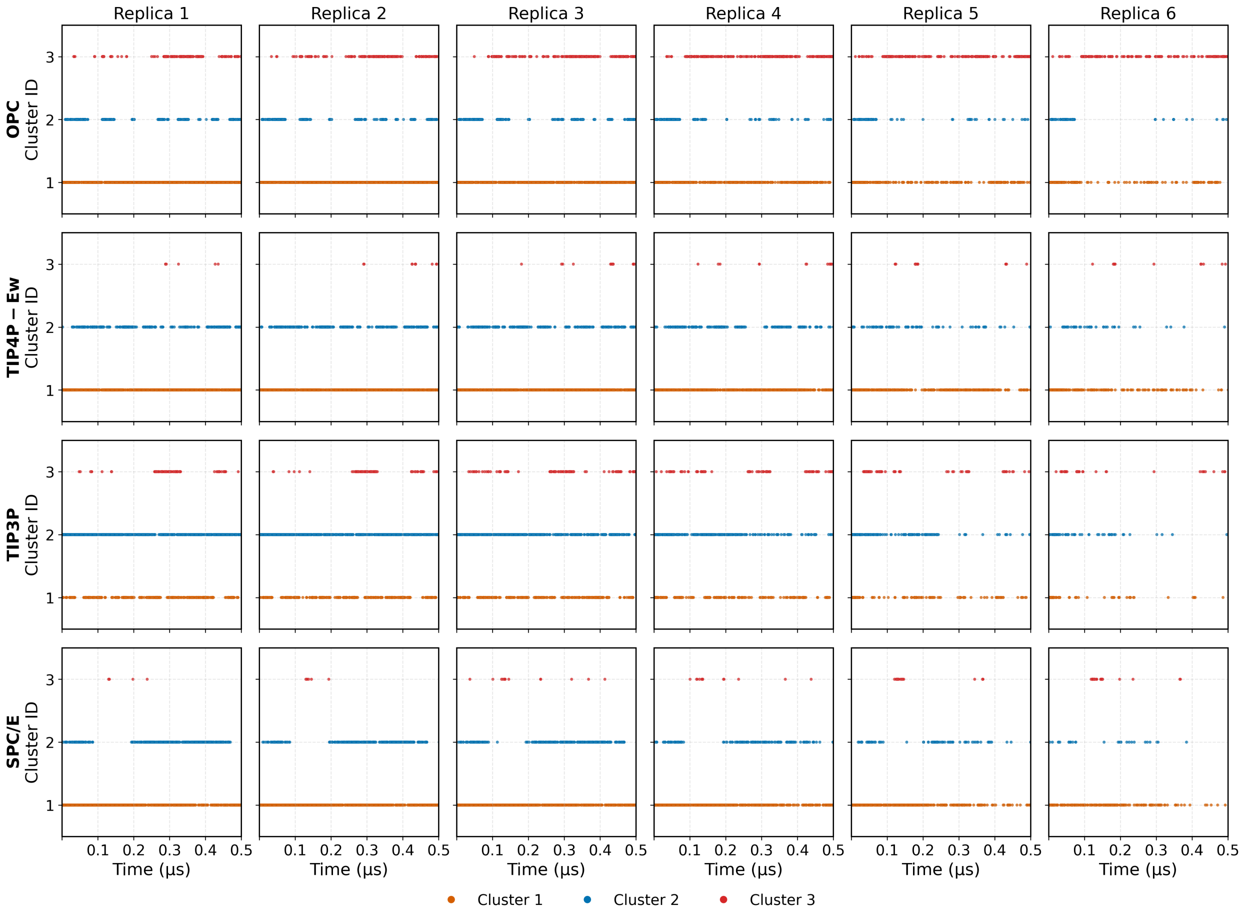


Figure S17: Time-resolved cluster trajectories of replicas across the four water models (OPC, TIP4P-Ew, TIP3P, and SPC/E). The clusters are color-coded as follows: Cluster 1 (orange), Cluster 2 (blue), and Cluster 3 (red).

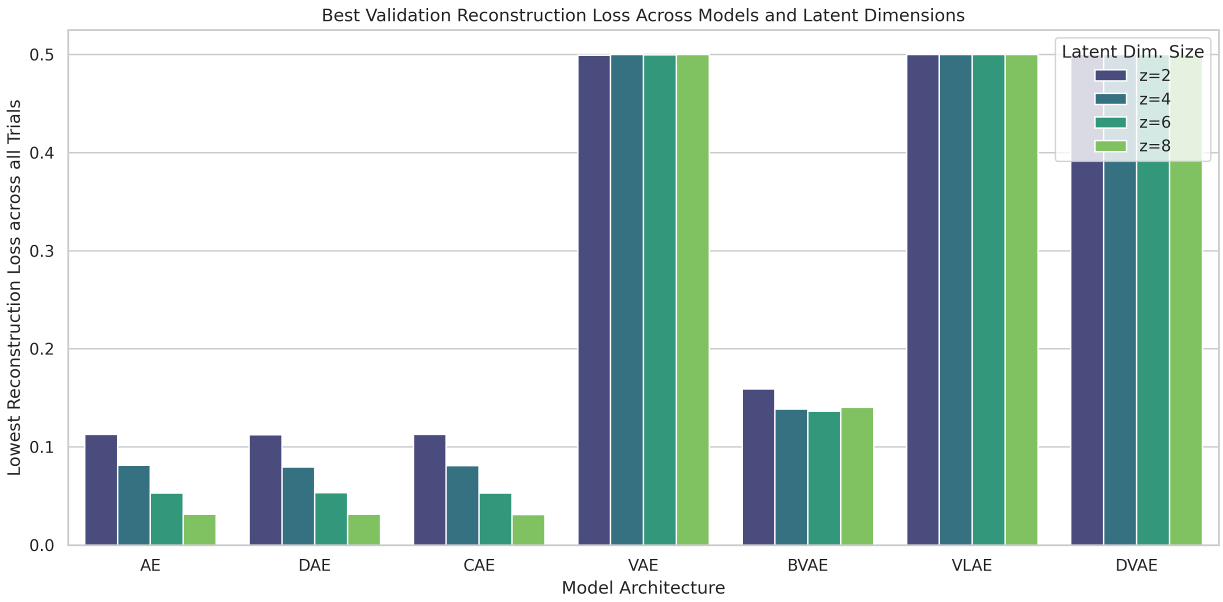


Figure S18: Reconstruction loss of AE, DAE, CAE, VAE, BVAE, VLAE, and DVAE across the corresponding embedding dimensions ($d = 2, 4, 6, 8$).
